# Cross-modal sensory compensation increases mosquito attraction to humans

**DOI:** 10.1101/2023.10.10.561721

**Authors:** Takeshi Morita, Nia G. Lyn, Ricarda K. von Heynitz, Olivia V. Goldman, Trevor R. Sorrells, Matthew DeGennaro, Benjamin J. Matthews, Leah Houri-Zeevi, Leslie B. Vosshall

## Abstract

Sensory compensation is a process that allows individuals with a loss of one sense, for instance hearing or vision, to adapt to changes in their sensory abilities. Where this phenomenon has been observed, there is enhanced perception by another sense to compensate for deficiency of the lost sense. Such compensation is important for humans and non-human animals that use multisensory integration for effective navigation and the execution of vital tasks. Among these, female mosquitoes are sensory specialists that rely heavily on integrating multiple human-emitted cues in their quest for a suitable host to obtain a blood meal. Here, we identify a previously undescribed mechanism of sensory compensation in female *Aedes aegypti* mosquitoes. Mutant mosquitoes lacking the odorant receptor co-receptor *Orco* show specific enhancement in heat-seeking behavior. This s compensation does not require the antenna, which was previously assumed to be the primary mosquito thermosensitive organ. Instead, we found that the tips of the forelegs are required to detect heat, and that the heightened sensitivity in heat detection is mediated by increased neuronal activity in foreleg sensory neurons, which are distant from the head appendage neurons that express *Orco*. By comparative gene expression analysis in wildtype and *Orco* mutant legs, we identify *Ir140*, a foreleg-enriched member of the Ionotropic Receptor (IR) superfamily of sensory receptors, as strongly upregulated in *Orco* mutant legs. Emphasizing the important role of IRs in thermosensation, we find that mutant mosquitoes lacking the IR co-receptor, *Ir25a*, lose all responses to heat, and *Ir140* mutants show strong deficits in responding to human skin temperatures. We generated an *Ir140*, *Orco* double mutant and show that these animals lose the remarkable sensory compensation seen in *Orco* mutants. This strongly suggests that upregulation of *Ir140* in the foreleg is the mechanism of sensory compensation in *Orco* mutants. Odorant receptor expression is sparse in legs, suggesting an indirect, long-range mechanism of sensory compensation. Our findings reveal a novel compensatory mechanism in which loss of one sensory modality in female *Aedes aegypti* mosquitoes results in greater sensitivity in another to maintain the overall effectiveness of their host-seeking behavior, further enhancing their status as the most dangerous predator of humans.

## INTRODUCTION

Animals are endowed with diverse sensory modalities to take in information from their environment, encompassing thermal, chemical, auditory, mechanical, visual, and other cues. These senses work together to form a representation in the brain of the sensory richness of the external world to guide appropriate behaviors. Because of the importance of multi-sensory integration, animals have mechanisms to adapt when they experience a loss or impairment in one of their sensory modalities, either through acute injury, chronic disease, or a congenital loss from birth. In this way, loss of vision, hearing, or touch, leads to the development of a heightened acuity in the remaining senses to compensate for their deficit. This adaptive process involves the reorganization and reallocation of neural resources to amplify the function of intact sensory modalities. For example, blind individuals often exhibit superior auditory and tactile perception as they adapt to prioritize these senses to navigate their environment safely and effectively (*1, 2*). Similarly, people with hearing loss may develop a more acute sense of vision or touch to compensate for their auditory impairment (*3*). Sensory compensation highlights the power of neural plasticity and capacity to adapt to changing sensory inputs, ultimately enabling individuals to maintain high perceptual awareness and functionality despite sensory impairment. Sensory compensation is a phenomenon observed throughout the animal kingdom. However, the full extent of its prevalence, the precise underlying mechanisms, and how it relates to each animal’s final behavioral outputs are not fully understood.

Sensory compensation is best understood at a mechanistic level in mammals and it involves anatomical modifications and reorganization at the thalamic and primary sensory cortex levels. These changes expand and refine connections within neighboring primary sensory cortical areas. For example, classical experiments in the ferret visual system revealed that eliminating retinal axon projection in one hemisphere led to alternative terminal space for these axons within the auditory thalamus (*4*). These changes were subsequently reflected in changes in cortical representation (*5, 6*). Functional reorganization also occurs between primary sensory cortical regions from different sensory modalities. Braille reading by blind individuals activates the primary visual cortex, suggesting an expansion of somatosensory function into brain regions generally dedicated to visual processing (*7*). Moreover, in congenitally blind subjects, the occipital lobe, typically associated with visual processing, is activated during auditory localization tasks (*8*).

Sensory compensation has also been observed by modulating existing neuronal circuits without requiring major anatomical rewiring. For example, *Caenorhabditis elegans* nematodes deprived of their sense of touch exhibited enhanced olfactory-mediated behavioral performance (*9*). This enhancement was attributed to the strengthening of synaptic transmission in the olfactory circuit that was a consequence of reduction in neuropeptide signaling cause by impaired mechanosensory circuits (*9*). Similarly, studies in visually deprived rats have revealed the prominent strengthening of α-amino-3-hydroxy-5-methyl-4-isoxazolepropionic acid (AMPA) receptor-mediated synaptic transmission in pyramidal neurons of layer 2/3 somatosensory barrel cortex (*10, 11*). These changes depended on long-distance serotonin signaling originating from the raphe nuclei (*10, 11*). More recently, a study conducted in *Drosophila melanogaster* flies showed that loss of olfaction led to an enhanced sensitivity to detect sugar that was mediated through an elevated sugar response by the protocerebrum anterior medial dopaminergic neurons in the mushroom body (*12*). These anatomical and functional studies highlight the diverse mechanisms that each sensory system and organism employs while also emphasizing the shared necessity of the process of sensory compensation. For organisms that depend heavily on integrating various sensory cues, this adaptive process is critical for guiding vital behavioral outputs.

**Figure 1:**
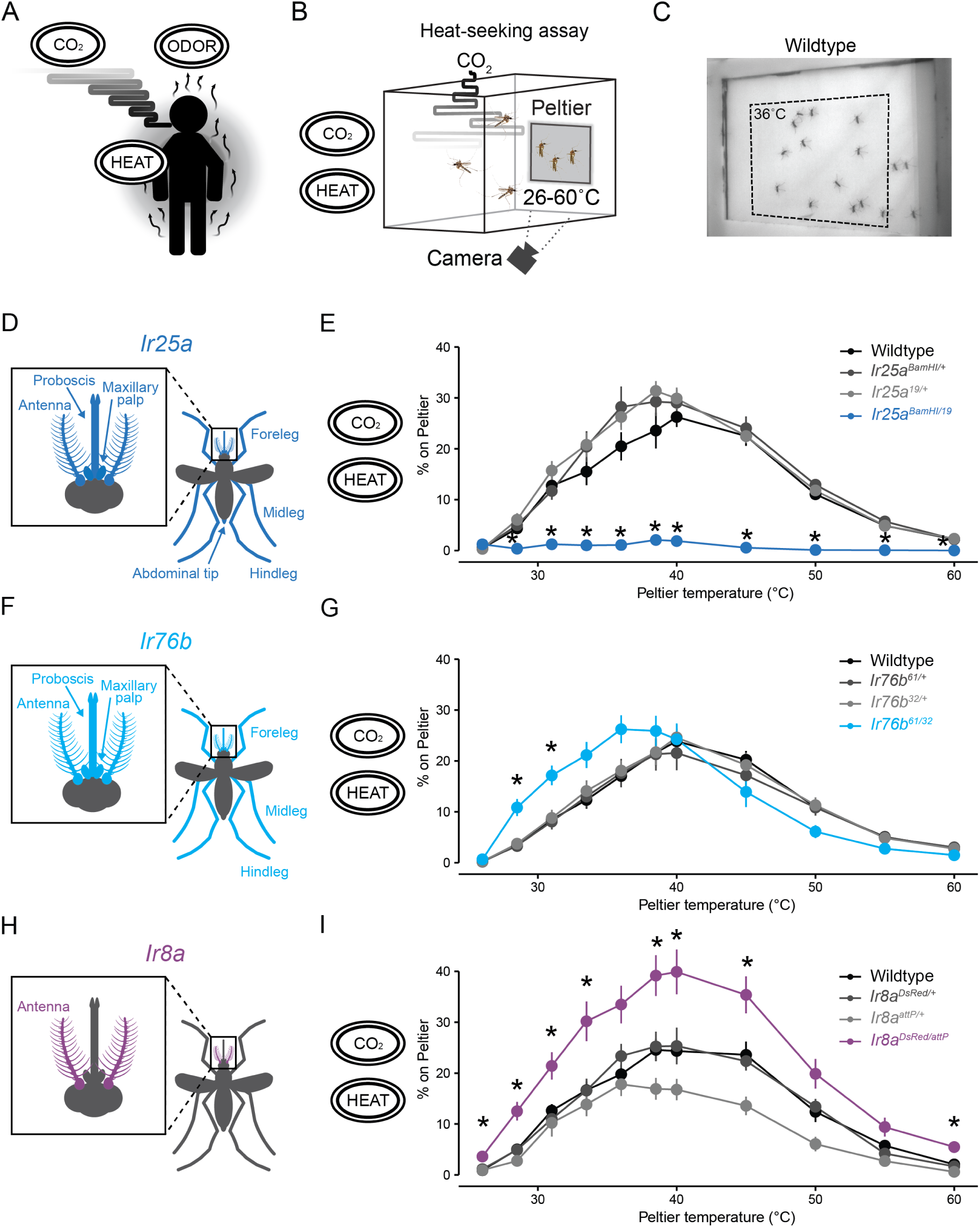
Ionotropic receptors tune thermotaxis behavior. (A) Female mosquitoes use redundant human-emitted cues (CO2, odor, and heat) to pursue blood meal. The black ovals with sensory cues displayed here and throughout the manuscript show the sensory stimuli present in the corresponding behavioral experiment. (B) Schematic of heat-seeking assay apparatus (30 cm^3^). (C) Representative experimental image of wildtype female mosquitoes on and near the Peltier (dotted line) set at 36°C. (D, F, H) Schematic of female body parts that express *Ir25a* (D), *Ir76b* (F), and *Ir8a* (H). (E, G, I) Percent of mosquitoes of indicated genotypes on Peltier during seconds 90-180 of stimuli of indicated temperature (mean ± SEM, n = 5-13 trials per genotype; datapoints marked with * indicate that the knockout mutant differs significantly from all other tested genotypes within each tested temperature at p < 0.05; one-way ANOVA with Tukey’s HSD post hoc test).

Mosquitoes are sensory specialists that detect and integrate diverse host-emitted cues, most notably, human-emitted odors, carbon dioxide (CO_2_), and body heat (Figure 1A). Unlike many other organisms, female mosquitoes need substantial blood meals to initiate egg production and can feed multiple times during their lifetime. This characteristic makes *Aedes aegypti* mosquitoes highly effective vectors for transmitting arboviruses such as dengue, yellow fever, Zika, and chikungunya, and makes *Anopheles gambiae* mosquitoes extremely dangerous vectors of the *Plasmodium* malaria parasites. Mosquitoes rely on three large multigene families to detect human-emitted cues, gustatory receptors (GRs), odorant receptors (ORs), and ionotropic receptors (IRs). These proteins form multi-subunit ligand-gated ion channel complexes featuring one or more ligand-selective subunit and an obligatory co-receptor subunit. While GRs are generally used for taste cues, members of this gene family form a heteromultimeric receptor that detects carbon dioxide (CO_2_) (*13–15*). In *Aedes aegypti*, there are 72 GRs, but for this gene family, the identity of the co-receptors is less well understood.Co-receptors for the OR and IR gene family are better understood and are central to their function. Each insect species can possess hundreds of ligand-selective ORs and IRs but only one odorant receptor co-receptor (Orco) and three IR co-receptors (*Ir8a*, *Ir76b*, and *Ir25a*) (*16*). The co-receptor subunit is required for the assembly and trafficking of functional ion channel complexes to the plasma membrane, and mutations of Orco and individual IR co-receptor subunits in *Drosophila melanogaster* disrupts the assembly of functional receptors (*16, 17*). In the case of *Aedes aegypti*, there are 116 ligand-selective ORs and 132 ligand-selective IRs (*18*). These expansive gene families of chemosensory receptors enable mosquitoes to detect various human-emitted cues. ORs generally respond to esters, alcohols, ketones, and aldehydes, while IRs have very flexible ligand tuning and have been shown to detect acids, amines, and physical stimuli, such as humidity and temperature (*19*). Due to this co-receptor configuration, the mutation of a single co-receptor gene can significantly impair an insect’s ability to detect entire classes of human-emitted sensory cues.

With the advent of genome engineering in this non-model organism (*20, 21*), it has been possible to show that genetic perturbations that disrupt each sensory modality in isolation have modest effects on mosquito attraction to humans. Mosquitoes lacking the CO_2_ receptor *Gr3* continue to exhibit attraction to humans under semi-field conditions (*13*). Mosquitoes with a loss-of-function mutation in *Orco* retain attraction to humans while losing a strong preference for humans over non-human animals, as well as becoming insensitive to the volatile effects of the insect repellent DEET (N,N-diethyl-meta-toluamide) (*20*). *Ir8a* mutants exhibit severe deficiencies in detecting lactic acid, a major component of human skin odor, but retain partial attraction to humans (*22*). *Ir76b* mutants remain attracted to humans despite losing their ability to feed on blood (*23*). *Ir21a* and *Ir93b* mutants experience reduced heat and humidity-seeking behavior but maintain their overall attraction to humans (*24, 25*). The recent discovery of extensive co-expression of ORs and IRs in single sensory neurons may explain this functional redundancy (*26, 27*). In addition, out of many sensory cues emitted by humans, detection of two cues is sufficient to initiate and synergize the drive for human-seeking behavior (*13*).

In this study, we discovered and investigated cross-modal sensory compensation between olfaction and thermosensation in the context of human-seeking behavior. We discovered that mutating the three IR co-receptors leads to different effects on thermotaxis behavior, ranging from complete loss of heat seeking (*Ir25a*), to a shift in preference to lower temperatures (*Ir76b*), or enhancement in sensitivity to heat (*Ir8a*). *Orco* mutant mosquitoes, which lack a functional OR pathway, show a remarkable increase in attraction to humans, which was fully attributable to their heightened sensitivity to heat. Contrary to our current understanding of insect thermosensation that points to the antenna as the major heat-sensitive organ, we discovered that mosquitoes employ their forelegs as an essential sensory structure for heat detection. The heightened heat-seeking behavior in *Orco* mutants can be traced back to enhanced thermosensitivity in these foreleg neurons. Comparative gene expression studies between wildtype and Orco mutant legs showed that both *Ir25a* and the ligand-specific IR subunit *Ir140* were upregulated in *Orco* mutant legs. Finally, we determined that *Ir140* plays a significant role in general heat-seeking behavior and that the enhanced thermosensitivity in *Orco* mutants is lost in *Ir140* mutants. Our results strongly support the notion that mosquitoes possess the capability for cross-modal sensory compensation, and that this compensation is due to long-range communication between head appendages and the foreleg. This novel mechanism allows these dangerous vector insects to maintain their overall effectiveness in human-seeking behavior even when one sensory modality is compromised.

## RESULTS

### Ionotropic receptors tune mosquito thermotaxis behavior

Our work began with the aim to identify genes involved in mosquito thermal attraction at the specific thermal range of human skin temperature (34_o_C-37_o_C). Our search used a previously described assay to model heat-seeking behavior by monitoring mosquitoes landing on a warmed Peltier element in a cage supplemented with CO_2_ (*13, 28*). This system can measure the attraction of mosquitoes to thermal stimuli by heating the Peltier element to a temperature ranging from ambient (26°C) to noxious (60°C) temperatures, and quantifying landing events. (Figure 1B, C, Supplemental Video S1). Using this system, we took a candidate gene approach by measuring thermotaxis behavior of mosquitoes carrying loss-of-function mutations in the three major IR co-receptors, *Ir25a*, *Ir76b*, *Ir8a*.

First, we looked at a broadly expressed IR co-receptor subunit, *Ir25a* (Figure 1D) (*29*). *Ir25a* is required for detecting acids, amines, humidity, cooling, and temperature-synchronized circadian clock oscillation in *Drosophila melanogaster* (*16, 30–34*). *Ir25a* also mediates amine detection and human odor attraction in *Aedes aegypti* (*26, 35*). The role of *Ir25a* in thermotaxis behavior in any mosquito species has not been examined. Using the heat-seeking assay, we found that *Ir25a* mutants failed to locate the heated Peltier element at all temperatures tested (Figure 1E). *Ir25a* mutants display normal CO_2_-evoked flight activity and noxious heat detection, suggesting that the failure to respond to heat is not due to locomotor deficits or a lack of behavioral participation (Supplemental Figure S1). These results indicate that *Ir25a* is required to detect all physiologically relevant temperatures. We next looked at another broadly expressed IR co-receptor subunit, *Ir76b* (Figure 1F). *Ir76b* is required for the detection of pH, amino acids, fatty acids, and salts in *Drosophila melanogaster* (*36–39*) and mediates amine detection and blood-feeding behavior in *Anopheles coluzzii* (*23*). The role of *Ir76b* in *Aedes aegypti* thermosensation has not been examined. We tested *Ir76b* mutants using the heat-seeking assay and found that these mutants showed normal responses to temperatures above 40°C but were more sensitive at lower temperatures (Figure 1G). This effect was specific to lower temperatures between 28.5°C and 31°C. These results suggest that *Ir76b* may be required to tune *Aedes aegypti* temperature preference towards human skin temperature. Finally, we looked at the antenna-enriched IR co-receptor subunit, *Ir8a*, which is required for lactic acid detection in *Drosophila melanogaster* and *Aedes aegypti* (*16, 22, 40*) (Figure 1H). Unlike *Ir25a* and *Ir76b* mutants, *Ir8a* mutants displayed enhanced heat-seeking behavior broadly across most temperature ranges tested (Figure 1I). Our analysis of heat-seeking behavior in these three IR co-receptor subunit mutants revealed distinct thermotaxis patterns for each mutant, implying that each IR co-receptor subunit contributes uniquely to the precise regulation of heat-seeking behavior in *Aedes aegypti*.

### *Orco* mutants display enhanced heat-seeking behavior

We next asked whether the Orco co-receptor is required for heat-seeking behavior in *Aedes aegypti*. In *Aedes aegypti*, *Orco* is mainly expressed in the antenna, proboscis, and maxillary palp (Figure 2A) (*20*). We tested *Orco* mutants using the heat-seeking assay to determine if ORs, like the IRs, had any signs of tuning the overall heat-seeking behavior in *Aedes aegypti*. Using the heat-seeking assay, we found that *Orco* mutants displayed enhanced heat-seeking behavior compared to their genetic controls at 36°C (Figure 2B, Supplemental Video S2). This enhancement was sustained during the entire duration of the heat stimulus period (Figure 2C). Furthermore, *Orco* mutants showed enhanced heat-seeking behavior, specifically towards human skin temperatures, and displayed normal avoidance behavior at the noxious temperature range (Figure 2B, D).

**Figure 2:**
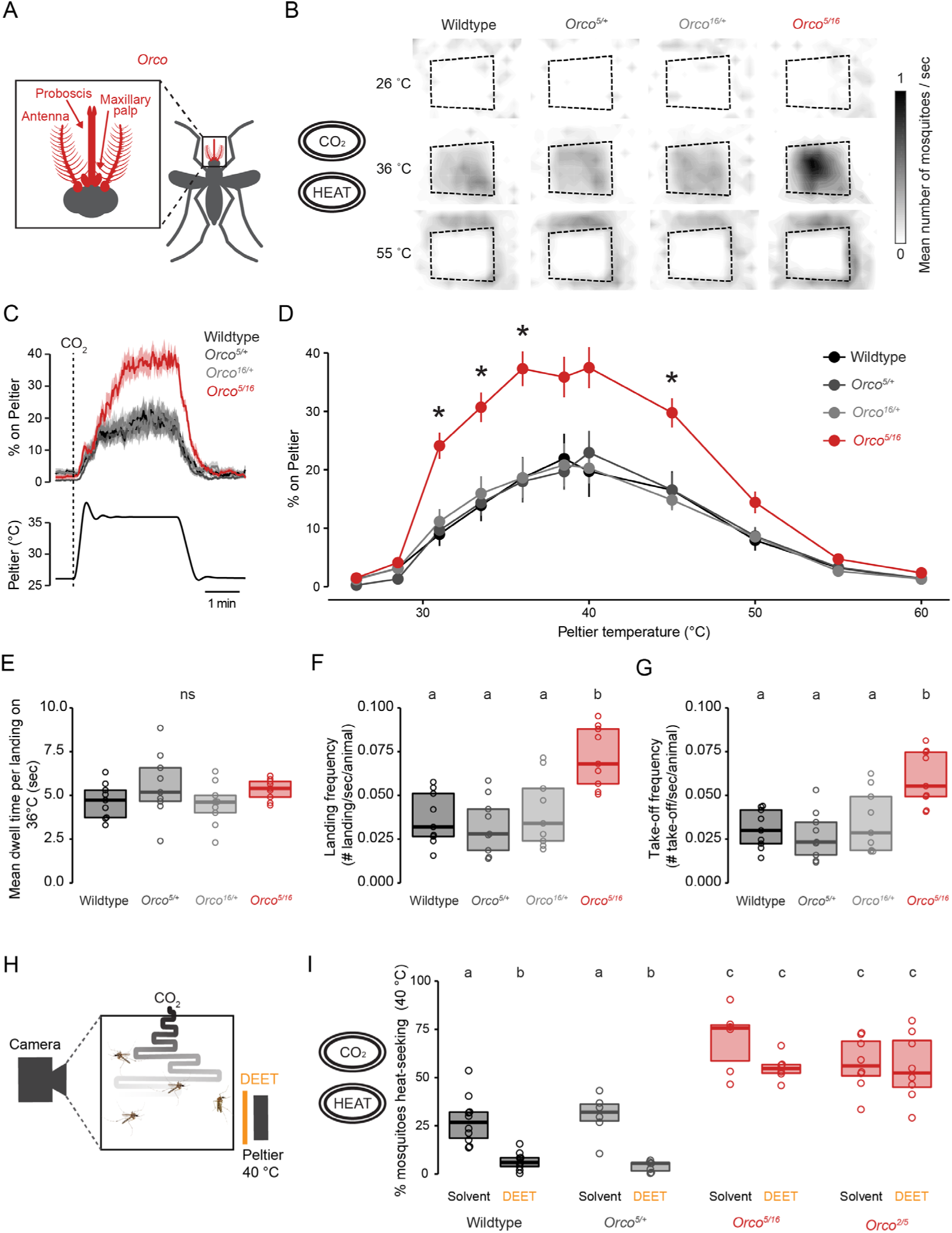
*Orco* mutant mosquitoes display enhanced heat-seeking behavior. (A) Schematic of female body parts that express *Orco*. (B) Heat maps showing mean mosquito occupancy for the indicated genotypes on the Peltier (dotted lines) and surrounding area at indicated Peltier temperature during seconds 90-180 of each stimulus period. (C) Mean ± SEM percent of mosquitoes of indicated genotypes on Peltier (top) during the 36°C trial (bottom). A 20 second pulse of CO_2_ was applied at the beginning of each stimulus period. (D) Percent of mosquitoes of indicated genotypes on Peltier during seconds 90-180 of stimuli of indicated temperature (mean ± SEM, n = 9 trials per genotype; datapoints marked with * indicate that the knockout mutant differs significantly from all other tested genotypes within each tested temperature at p < 0.05; one-way ANOVA with Tukey’s HSD post hoc test). (E-G) Mean dwell time (E), landing frequency (F), and take-off frequency (G) of indicated genotypes on the Peltier surface during the 36°C trial (n = 9 trials/genotype). (H) Schematic representation of the modified heat-seeking assay (28 cm^3^) in the presence of DEET. (I) Percent of mosquitoes of indicated genotype and treatment groups heat-seeking during a 40°C trial (n = 6-10 trials/genotype). Data are plotted as scatter-box plots (individual data points, median as horizontal line, interquartile range as box) for (E,F,G,I). Data labeled with different letters differ significantly (p < 0.05; one-way ANOVA with Tukey’s HSD post hoc test at each tested temperature).

Next, we investigated which aspect of mosquito heat-seeking behavior was specifically altered in *Orco* mutants. We first tested whether *Orco* mutants spent longer on the heated surface by calculating the average dwell time per landing event for each animal at human skin temperature. Our analysis revealed that *Orco* mutants spent equal time as controls on the Peltier element per landing event (Figure 2E). Another possibility was that *Orco* mosquitoes engaged in landing events more frequently than their genetic controls. We analyzed the final 60 seconds of each stimulus period and calculated both landing and take-off event frequencies for each animal. We found that *Orco* mutants display increased landing and take-off frequencies, demonstrating that *Orco* mutants had persistent and sustained engagement towards the heated Peltier element throughout the heat trials (Figure 2F, G).

Given the dramatic increase in the attraction of *Orco* mutants to heat, we next asked if the insect repellent DEET deters these mutants from heat-seeking behavior. Unlike wildtype animals, *Orco* mutants are attracted to human odor in the presence of DEET, however whether volatile DEET interferes with heat-seeking independently is not known (*20*). We designed a modified version of the heat-seeking assay by placing a source of DEET in front of the Peltier device set at 40°C (Figure 2H). This source of DEET was placed across a spacer to avoid direct contact of DEET contact by the mosquitoes. Contact chemorepellency is mediated by the taste system of the mosquito leg, and does not require *Orco* function (*41*). As expected, we found that DEET was sufficient to drive mosquitoes away from heat in wildtype and heterozygous controls. However, *Orco* mutant mosquitoes retained their enhanced heat-seeking behavior even in the presence of DEET (Figure 2I). We replicated this result with another heteroallelic mutant, *Orco_2/5_* (Figure 2I). These data indicate that enhanced heat-seeking is unaffected by *Orco*-mediated repellency, thus depending on the absence of *Orco*. Additionally, these data demonstrate the critical role of *Orco* in DEET detection and avoidance behavior in the presence of multi-modal human-related cues (*20*).

### *Orco* mutants display enhanced human-seeking behavior

We next investigated sensory cues of different modalities to test the specificity of the enhanced thermotaxis behavior displayed by *Orco* mutants. Using the CO_2_ activation assay, we tested the activity and arousal of *Orco* mutants in response to a brief 20-second pulse of CO_2_ (Figure 3A, Supplemental Figure S2A). CO_2_-evoked flight activity and total distance traveled in response to CO_2_ were indistinguishable from genetic controls (Figure 3B, Supplemental Figure S2B,C). We further tested whether attraction toward human odor was altered in *Orco* mutants using the nylon-next-to-cage assay (Figure 3C, E). A previous study showed that *Orco* mutants show normal attraction to humans but are impaired in discriminating human from non-human odors (*20*). Consistent with these previous observations, *Orco* mutants show normal attraction towards human-worn nylon (Figure 3E, F). This attraction was specific to host odors because mosquitoes were not attracted to unworn nylon (Figure 3C, D). We further tested whether the enhanced thermosensitivity of *Orco* mutants enhances their ability to find live human hosts. To test this, we designed an arm-next-to-cage assay with a human arm positioned close to the cage to allow mosquitoes to detect human-emitted odor, CO_2,_ and body temperature without physical contact (Figure 3G). We found that *Orco* mutants displayed enhanced human arm-seeking behavior compared to genetic controls (Figure 3H). We further repeated the experiment in the presence of volatile DEET to test whether the enhanced human-seeking behavior is retained in the presence of this repellent (Figure 3I). We found that *Orco* mutants were still capable of human arm-seeking behavior in the presence of DEET (Figure 3J). These data, along with the heat-seeking data in the presence of DEET (Figure 2I), suggest that enhancement of *Orco* human-seeking behavior is unaffected by DEET.

**Figure 3:**
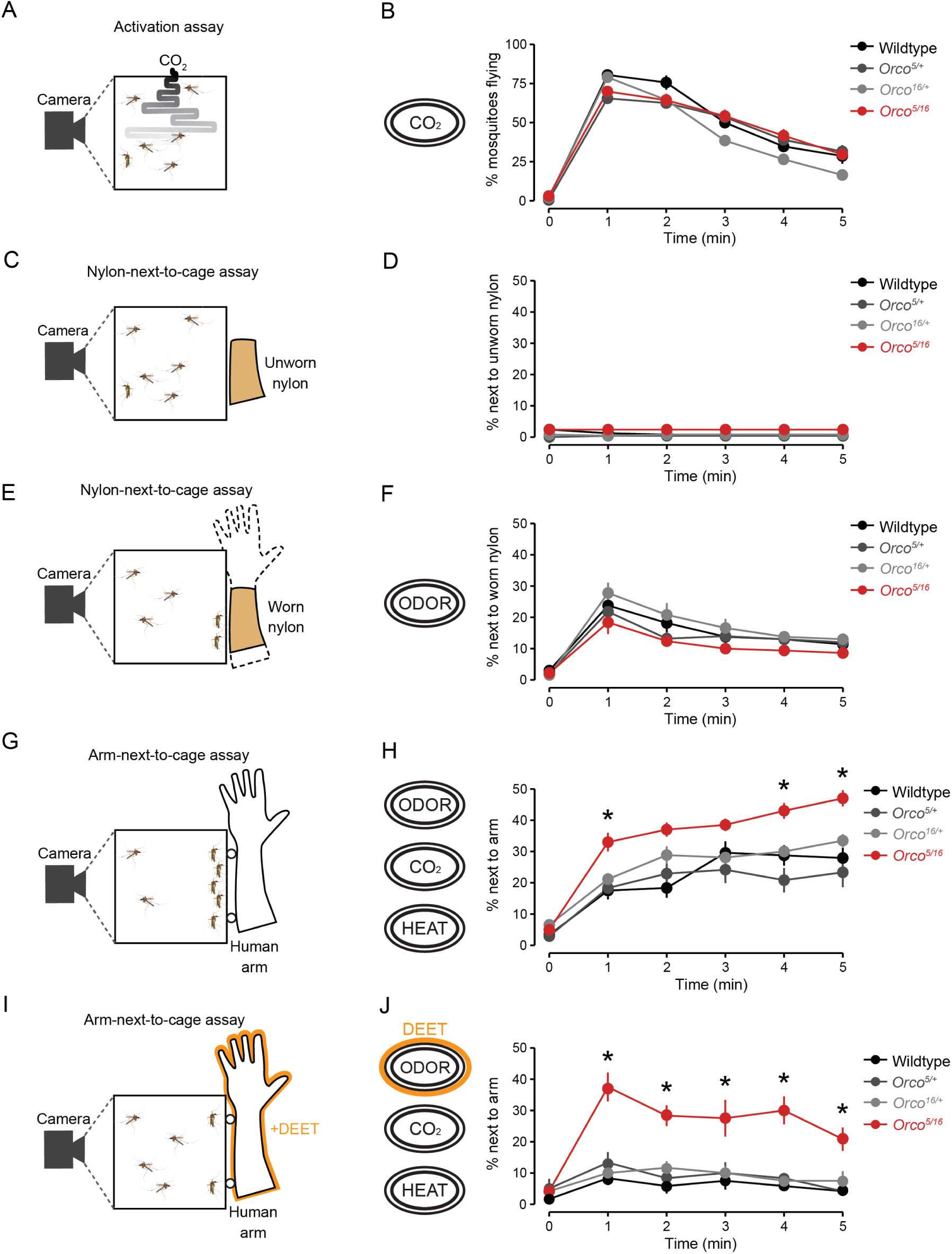
Human attraction is enhanced in *Orco* mutant mosquitoes. (A) Schematic of the CO_2_ activation assay. (B) Percent mosquitoes flying in the presence of CO_2_ quantified every minute (mean ± SEM, n = 10 trials per genotype). (C, E) Nylon-next-to-cage assay schematic with unworn (C) and worn (E) nylon. (D, F) Percent mosquitoes next to unworn (D, n=10 trials/genotype) or worn (F, n=5 trials/genotype) nylon, quantified once per minute (mean ± SEM). (G, I) Arm-next-to-cage assay schematic with human arm without (G) or with 10% DEET (I). (H, J) Percent mosquitoes next to the human arm without (H, n=10-13 trials/genotype) or with 10% DEET (J, n= 6 trials/genotype) quantified once per minute (mean ± SEM). In panels D, F, H, and J, datapoints marked with * indicate that knockout mutant differs significantly from all other tested genotypes within each tested temperature at p < 0.05; one-way ANOVA with Tukey’s HSD post hoc test).

### Forelegs mediate thermotaxis behavior

Previous studies in *Drosophila melanogaster* have identified thermosensitive sensory structures and neurons activated by diverse thermoreceptors and ion channels (Figure 4A) (*31, 34, 42–46*). In mosquitoes, prior studies identified a set of thermosensitive neurons at the most distal antennal segment that responded to cooling or heating air (Figure 4B, C) (*24, 25, 47–49*). Since *Orco* is expressed in the antenna, we hypothesized that loss of *Orco* may directly affect the function of thermosensitive neurons at the antennal tip. However, none of the previous studies investigated the direct roles and requirements of these thermosensitive neurons in the context of thermotaxis behaviors. To test this, we removed the three most distal antenna segments by cutting off the tip and performed thermotaxis assays (Figure 4D). We first wanted to confirm that antennal tip removal had minimal consequences on the overall behavior of the mosquitoes using an arm-feeding assay where we measured the ability of the antenna tip-cut mosquitoes to blood-feed on a human arm (Figure 4E). Antenna tip-cut mosquitoes engorged on human arms comparable to fully intact animals (Figure 4F). This result confirms that the antenna tip removal had minimal effect on overall human-seeking behavior, consistent with previous reports (*50, 51*).

**Figure 4:**
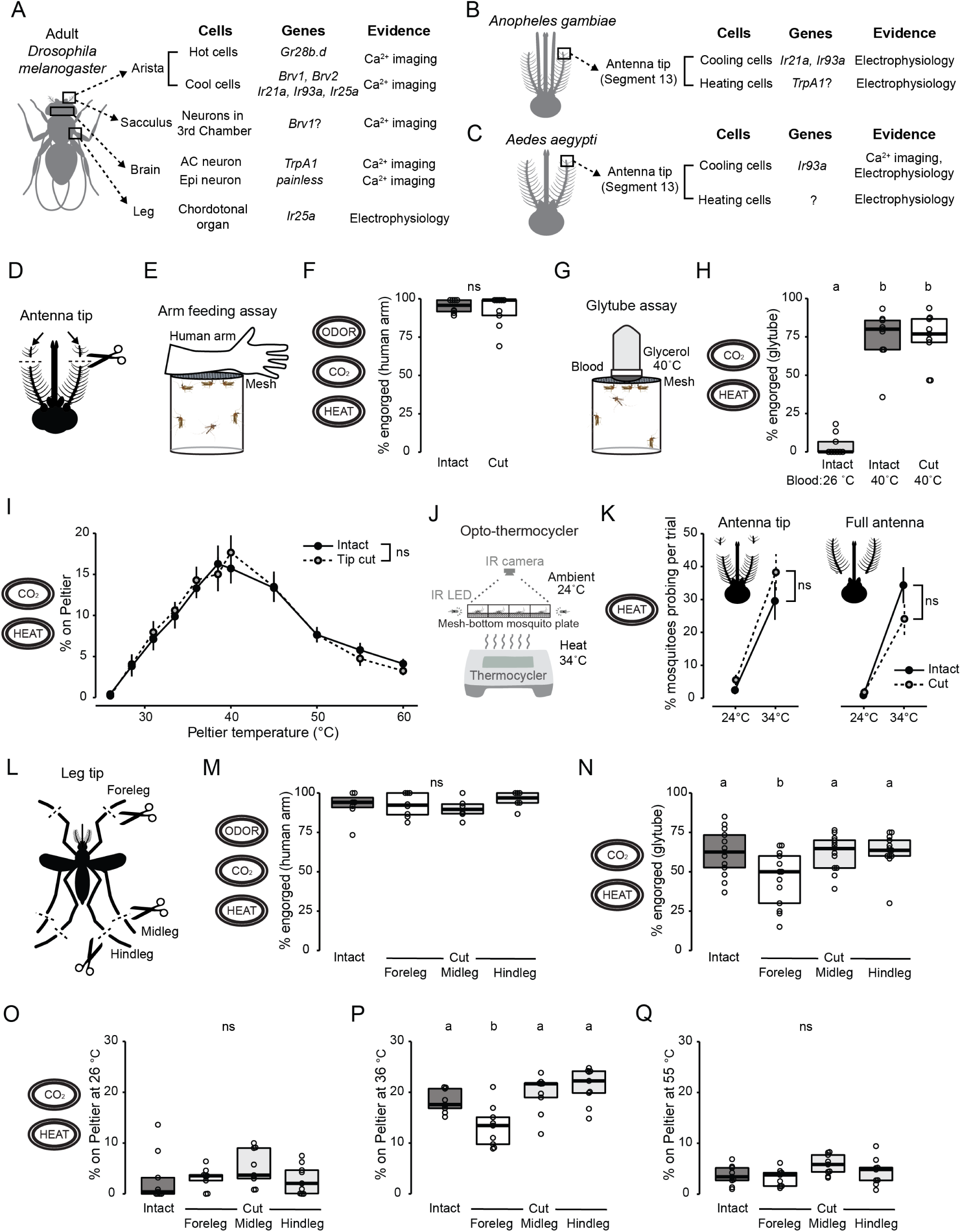
Forelegs but not antennae mediate *Aedes aegypti* heat-seeking behavior. (A-C) Sensory organs, genes, and experimental evidence for intrinsically thermosensitive neurons currently known in adult *Drosophila melanogaster* (*31, 34, 42–46*) (A), *Anopheles gambiae* (*24, 25, 49*) (B), and *Aedes aegypti* (*25, 47, 48*) (C). (D) Schematic of antennal tip removal, with distal three segments cut from both antennae. (E) Arm feeding assay schematic. (F) Percent engorged on the human arm of indicated antenna treatment (n = 8-9 trials/genotype). (G) Glytube assay schematic. (H) Percent engorged on blood in the Glytube assay of indicated antenna treatment and blood temperature (n = 9 trials/condition). (I) Percent of mosquitoes of indicated antenna treatment on Peltier during seconds 90-180 of stimuli of indicated temperature (mean ± SEM, n = 8-9 trials per treatment; ns, unpaired t-test at each tested temperature). (J) Schematic of the Opto-thermocycler assay. (K) Percent probing in response to ambient temperature and heat at 34°C for antennal tip removal (left, n = 9 trials/condition) and full antenna removal (right, n = 8 trials/condition). Data are plotted as mean ± SEM (ns, unpaired t-test comparing cut and intact mosquitoes at 34°C). (L) Tarsal removal schematic, with distal three tarsal segments cut from each pair of legs. (M) Percent engorged on the human arm of indicated tarsal treatment (n = 6-8 trials/condition). (N) Percent engorged on blood in the Glytube assay of indicated tarsal treatment (n = 13 trials/condition). (O-Q) Percent of mosquitoes of indicated genotypes on Peltier during the 26°C (O), 36°C (P), or 55°C (Q) trials. n= 9 trials/genotype. Data are plotted as scatter-box plots [individual data points, median as horizontal line, interquartile range as box) for (F,H,M-Q)]. Data labeled with different letters are significantly different (p < 0.05; one-way ANOVA with Tukey’s HSD post hoc test).

We then investigated the role of antenna tip neurons in thermotaxis behavior, first using a Glytube assay, an artificial blood-feeding system that provides mosquitoes with warm blood in the presence of CO_2_ (Figure 4G) (*52*). Surprisingly, we found that antenna tip-cut animals still engorged on blood at comparable levels to fully intact mock-treated animals (Figure 4H). We then turned to the heat-seeking assay to monitor heat-seeking behavior during each heat stimulus. Consistent with Glytube assay results, antenna-tip-cut mosquitoes displayed normal heat-seeking behavior, indistinguishable from fully intact mock-treated controls at all temperatures tested (Figure 4I). Because our data show that the first three segments of the antennal tip are dispensable for various thermotaxis tasks, we wanted to know whether any portion of the antenna was required to detect thermal information. Since complete antenna removal disables mosquitoes from flying (*50, 51*), we turned to the opto-thermocycler assay (53). This assay tracks thermotaxis behavior, using probing movement as a proxy for heat detection at high temporal and spatial resolution without requiring flying behavior (Figure 4J). The antenna tip-cut animals displayed probing behavior indistinguishable from mock-treated controls, consistent with our other thermotaxis assays. Furthermore, animals with the entire antenna removed also detected and responded by displaying probing behavior to a thermal stimulus indistinguishable from fully intact mock-treated controls (Figure 4K). These experiments suggest that the antenna tip and even the entire antenna is dispensable for heat detection in *Aedes aegypti*.

We further tested the possibility that other sensory appendages are responsible for mosquito thermotaxis behavior. We focused on the legs because these sensory appendages make direct contact with heated surfaces, and a previous study showed that *Drosophila melanogaster* leg neurons responded directly to thermal stimuli (*31*). Ticks also have a unique sensory organ on their distal leg segment called Haller’s organ that functions as a heat sensor (54). To determine whether legs contribute to thermotaxis behavior in *Aedes aegypti*, we removed the three most distal segments from each pair of legs – forelegs, midlegs, and hindlegs (Figure 4L). Cutting the tips of each pair of legs had no significant effect on human-seeking and blood-feeding behavior on a live human arm (Figure 4M). However, animals with foreleg tips cut had a significant reduction in their ability to engorge on blood in the Glytube assay, where the only cues presented to these animals were heat and CO_2_ (Figure 4N). Furthermore, animals with foreleg tips cut had a significant reduction in heat-seeking behavior to the Peltier element set to 36_o_C (Figure 4P) but not at ambient (Figure 4O) or at a noxious temperature (Figure 4Q). These data suggest that mosquito forelegs significantly contribute to *Aedes aegypti* thermotaxis behavior.

### Neuroanatomy of the *Aedes aegypti* legs

The neuroanatomy of *Aedes aegypti* legs has not been extensively studied. Insect legs comprise multiple segments -coxa, trochanter, femur, tibia, and tarsus. We focused on the tarsus, explicitly looking at the fifth tarsomere (Figure 5A), which is the most distal tarsal segment and is considered the primary site for chemoreception in the leg (*55, 56*). We characterized the neuronal anatomy of the fifth tarsomere using a panel of chemosensory receptor driver lines, each expressing a fluorescent reporter, *dTomato,* in its corresponding neural population (Figure 5B-E). Specifically, we used *brp-QF2w > QUAS-dTomato* to label all neurons (*57*), *Ir25a-QF2 > QUAS-dTomato* and *Ir76b-QF2 > QUAS-dTomato* to label putative thermosensitive and chemosensitive cells (*26*), and *Gr4-QF2 > QUAS-dTomato* to label sugar-sensing neurons previously shown to induce appetitive behavior once activated (*58*). *Aedes aegypti* tarsi are densely covered with scales, and the neuronal cell bodies extended their dendrites into sensilla that decorate the leg.

**Figure 5:**
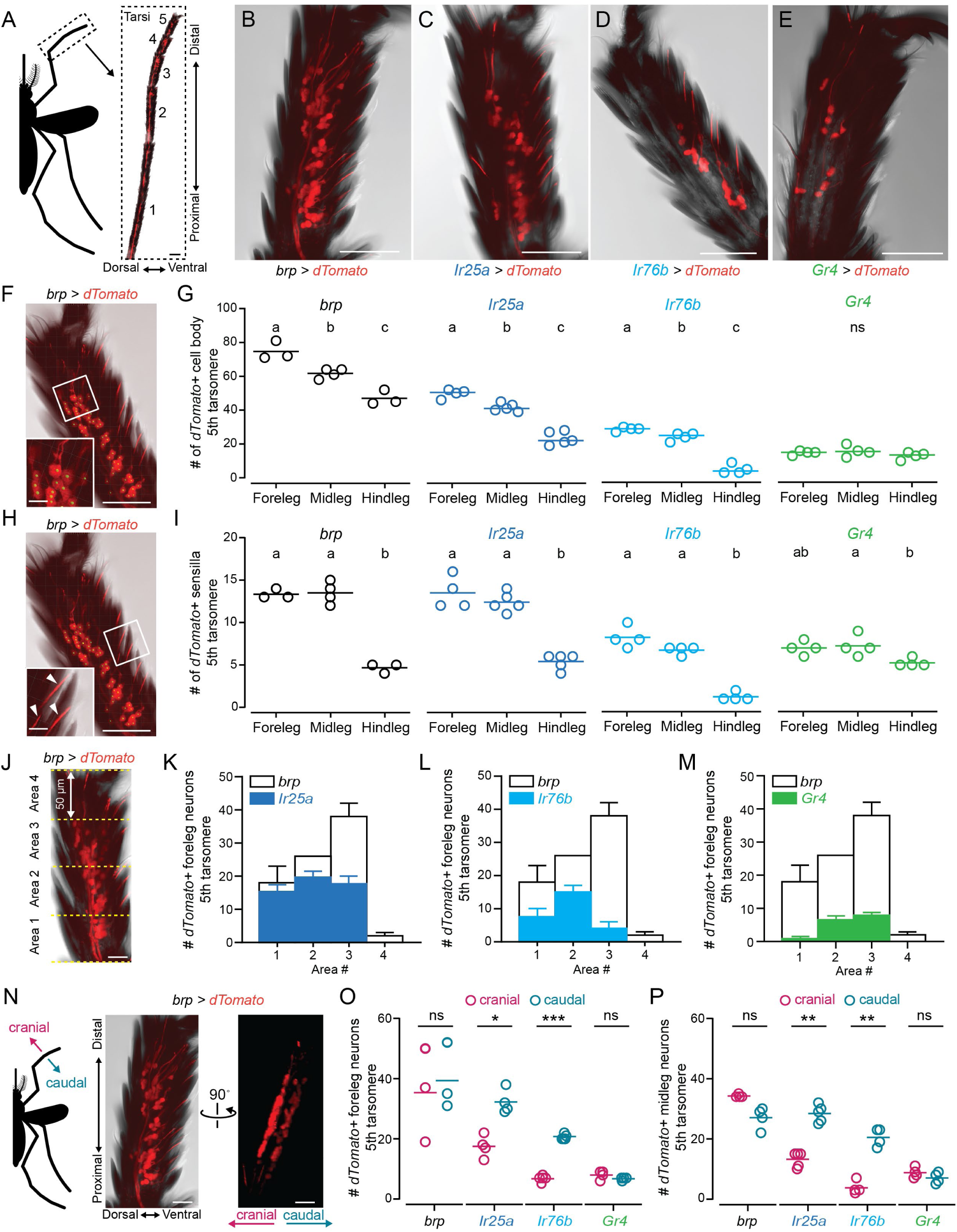
Morphology of tarsal neurons. (A) Schematic representation of a mosquito leg and representative tiled confocal image of tarsal segments of *brp*-*wQF2>QUAS-dTomato* mosquitoes expressing dTomato in all neurons. For this and all other images in the figure, dorsal is left and distal is top. Individual tarsomeres are numbered from proximal to distal segments.(B-E) Maximum-intensity projections of confocal z-stack images of dTomato expression in the fifth tarsomere of forelegs in *brp*-*wQF2>QUAS-dTomato* (B), *Ir25a-QF2>QUAS-dTomato* (C), *Ir76b-QF2>QUAS-dTomato* (D), and *Gr4-QF2>QUAS-dTomato* (E) animals. (F,H) Representative image of the fifth tarsomere used for cell body (F) and sensilla (H) quantification. (Inset) Zoomed-in image of the region outlined with a white box. Cell bodies are labeled with green dots and representative sensilla are labeled with white arrowheads. (G,I) Quantification of dTomato-labeled cell bodies (G) and sensilla (I) in the fifth tarsomere in the foreleg, midleg, and hindleg of indicated reporter lines (n = 3-5 tarsi). Data are plotted as scatter plots (individual data points, mean as horizontal line). Data labeled with different letters differ significantly within each reporter line (p < 0.05; one-way ANOVA with Tukey’s HSD post hoc test). (J-M) Distribution of dTomato-expressing neurons along the proximal-distal axis of the fifth tarsomere in forelegs from *Ir25a-QF2>QUAS-dTomato* (K), *Ir76b-QF2>QUAS-dTomato* (L), and *Gr4-QF2>QUAS-dTomato* (M). The fifth tarsomere is divided into four 50 µm areas indicated with yellow dotted lines (J). Bars outlined with black lines indicate *brp-*positive neurons in the same area from a different sample (n = 2-4 tarsi). (N) Schematic representation of the cranial-caudal axis and 3D projection of the fifth tarsomere shown in (B) rotated approximately 90 degrees. (O,P) Number of dTomato-labeled cell bodies in individual planes in the fifth tarsomere of forelegs (O) and midlegs (P) labeled by the indicated driver lines (n = 3-5 tarsi). Data are plotted as scatter plots (individual data points, mean as horizontal line; datapoints marked with *, **, or *** indicate statistically significantly distinguishable groups using paired t-test, p < 0.05, p < 0.01, p < 0.005, respectively. Scale bars = 100 µm for (A), 50 µm for (B-F,H,J,L), and 10 µm for (Insets in F,H).

Using the *brp-QF2w > QUAS-dTomato* reporter line, we quantified the number of neurons within the fifth tarsomere of each leg and observed a decreasing order of neuron count from forelegs to midlegs and hindlegs (Figure 5F,G). We identified an average of 75 total neurons in the fifth tarsomere of the foreleg, 62 in the fifth tarsomere of the midleg, and 47 in the fifth tarsomere of the hindleg (Figure 5B,G). We then examined reporter expression for *Ir25a*, *Ir76b*, and *Gr4*. *Ir25a-*expressing neurons were found in roughly two-thirds of the total fifth tarsal neurons, with a decreasing number in order of forelegs, midlegs, and hindlegs (Figure 5C,G). *Ir76b*-expression was less abundant, in one-third of the neurons of the forelegs and midlegs and, to a lesser extent, in the hindlegs (Figure 5D,G). In contrast, expression of the *Gr4* sugar receptor was equivalent in all three legs (Figure 5E,G).

In our images, we observed and quantified dendritic processes from cell bodies into their respective sensilla (Figure 5H). As with the total neural count, the total sensilla count (*brp-* positive) was much higher in the fifth tarsomere of forelegs and midlegs compared to hindlegs (Figure 5B, I). The number of *Ir25a*-positive sensilla count mirrored that of *brp*-positive sensilla count, suggesting that *Ir25a*-positive neurons innervate all or most chemosensory sensilla (Figure 5C, I). *Ir76b*-positive sensilla represented half of the total sensilla counts in all three legs (Figure 5D, I). The *Gr4*-positive sensilla count represented half of the total *brp*-positive sensilla counts in the forelegs and midlegs and had roughly the same number of hindleg sensilla (Figure 5E, I) (*26*).

We further investigated the spatial distribution of each chemoreceptor-expressing neuron within the fifth tarsomere. We divided the fifth tarsomere into 50 µm regions and counted the neurons in each region (Figure 5J). *Ir25a*-expressing neurons were broadly distributed across the entire fifth tarsomere while *Ir76b*-expressing neurons were more concentrated in the proximal tarsal region. *Gr4*-expressing neurons were primarily located in the distal area of the fifth tarsomere (Figure 5K-M).

We noticed the neurons were organized in two planes on each side of the tarsus (Figure 5N, Supplemental Video S3). To our knowledge, this has not been described in other insects. When the mosquito leg is outstretched, one side of the tarsus faces toward the head and the other side is oriented toward the caudal end of the abdomen. Thus, we named these two neural planes the “cranial” and “caudal” sides of the tarsus and quantified the number of labeled neurons per plane for each reporter line. While the overall number of foreleg neurons of the fifth tarsomere in each plane was similar, *Ir25a* and *Ir76b* represented a higher proportion of neurons on the caudal side compared to the cranial side, while *Gr4*-expressing neurons displayed an even distribution between the two sides (Figure 5O). This cranial-caudal distribution was similar in the midleg (Figure 5P).

### Mosquito leg neurons respond to heat

We next asked which neurons in each of the main *Aedes aegypti* sensory appendages responded to heat. We used the pan-neuronal imaging line *brp-QF2w > QUAS-dTomato-T2A-GCaMP6s* to measure Ca_2+_ response as a proxy for neuronal activity in response to heated air in the four main sensory appendages – antenna, maxillary palp, proboscis, and legs (Figure 6A). We developed a functional imaging setup that allowed us to acquire GCaMP and dTomato fluorescence signals while applying heated air over the samples. Dual channel imaging is critical to measuring heat-evoked Ca_2+_ activity due to the intrinsic thermosensitive properties of fluorescent proteins. We report the change in the ratio of the GCaMP signal over the dTomato signal (*R = GCaMP6s/*dTomato) normalized to baseline (*ΔR/R*) to correct for any movement artifact as well as temperature-evoked intrinsic changes in fluorescence signals as previously described to study thermosensitivity in *C. elegans* thermosensory neurons (*59, 60*) (Figure 6B, Supplemental Figure S3A-C).

**Figure 6:**
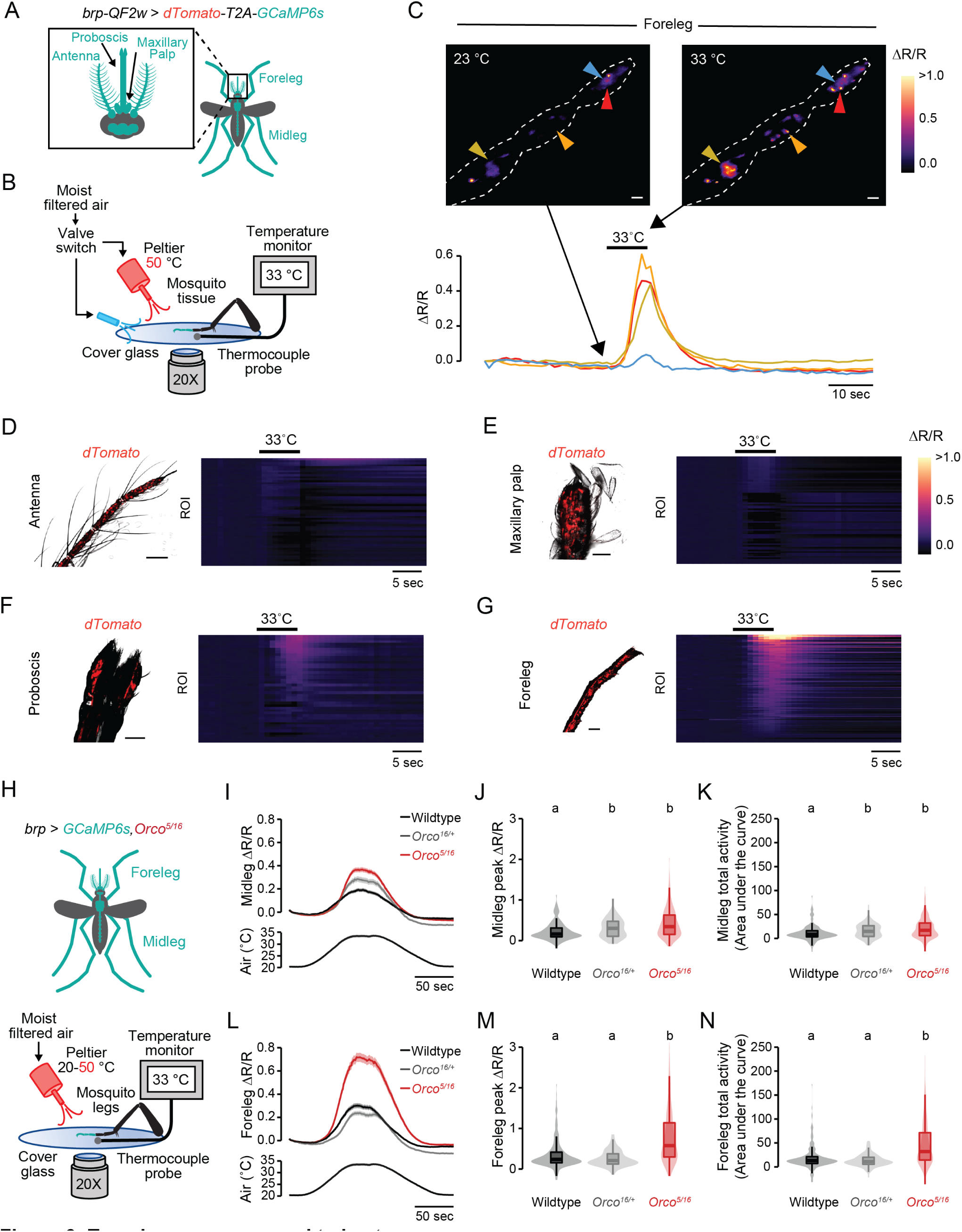
Tarsal neurons respond to heat. (A) Schematic of female body parts that express GCaMP6s in the pan-neuronal imaging strain *brp-QF2w > dTomato-T2A-GCaMP6s*. (B) Schematic of *ex vivo* ratiometric sensory appendage Ca^2+^ imaging setup. (C) Wide-field images of foreleg distal tarsal segments, prior to and during heat stimulus. Neuronal responses are indicated as pseudo-color images (top). Calcium traces from representative cells are indicated by the arrowheads of the same color in the top image (bottom). The black bar indicates the duration of the heat stimulus. (D-G) Wide-field dTomato image (left) and heatmap showing calcium imaging response for individual ROI across time (right) in the antenna (D, n = 54 ROI), maxillary palp (E, n = 54 ROI), proboscis (F, n = 42 ROI), and forelegs (G, n = 126 ROI). (H) Schematic of female body parts that express GCaMP6s in the pan-neuronal imaging strain *brp-QF2w > dTomato-T2A-GCaMP6s* in the *Orco^5/16^* mutant background (top) and schematic of *ex vivo* ratiometric sensory appendage Ca^2+^ imaging setup. (I-K, L-N) *Orco* mutant midleg (I-K) and foreleg (L-N) responses to warm air showing average Ca^2+^ responses across time (I, midleg; L, foreleg), peak responses (J, midleg; M, foreleg), and total activity (K, midleg; N, foreleg) of indicated genotypes (I-K: n = 91-265 total ROI from 4-10 trials/genotype; L-N: n = 102-260 total ROI from 4-10 trials/genotype). Data are plotted as mean ± SEM for (I,L) and violin-box plots (median as horizontal line, interquartile range as box, and 5th and 95th percentiles indicated as whiskers) for (J,K,M,N). Data labeled with different letters are significantly different (p<0.05, Kruskal-Wallis test with Dunn’s multiple comparisons). Scale bars = 50 µm for (C-G).

We first tested whether foreleg neurons responded to heated air. Consistent with our behavioral data, foreleg neurons throughout all three distal segments responded robustly to heated air (Figure 6C). These heat-evoked foreleg neural responses were reproducible within each trial (Supplemental Figure S3D). This response was independent of any mechanical artifact from the valve switching between two air outlets, because neuronal response returned after the heat source was turned back on (Supplemental Figure S3E, F). We then tested the specificity of this heat-evoked neuronal response in other sensory structures. We could not detect Ca_2+_ activity in the antenna or maxillary palp (Figure 6D, E), while the proboscis had weak responses to heated air (Figure 6F). By far, the most robust heat-evoked activity was seen in foreleg neurons (Figure 6G). These functional imaging data further suggest the critical role of foreleg neurons, and the dispensability of the antenna, during thermotaxis behavior.

Now that we identified the primary sensory appendage for heat detection, we tested whether the foreleg is the primary site for enhanced thermosensitivity in *Orco* mutants. To test this, we generated heteroallelic *Orco* mutant pan-neuronal imaging mosquitoes by crossing two different *Orco* mutant alleles into the *brp-QF2w > QUAS-dTomato-T2A-GCaMP6s* imaging background strain (Figure 6H). We modified the imaging setup to apply a slow heat ramp using a single Peltier outlet to capture any heat-evoked activation threshold differences in *Orco* mutant leg neurons. Using these *Orco* mutant pan-neuronal GCaMP mosquitoes and the modified imaging setup, we found that both heterozygous and *Orco* knockout mutant midlegs, although indistinguishable from each other, displayed higher peak response and total activity in response to heated air than wildtype controls (Figure 6I-K). This heterozygous effect was unexpected because *Orco* typically displays a fully recessive phenotype (*20*). In contrast, peak amplitude and total activity were enhanced in *Orco* knockout forelegs compared to both wildtype and heterozygous controls (Figure 6L-N). These data suggest that the enhanced thermosensitivity in the foreleg neurons underlies the increased thermotaxis behavior of *Orco* mutants.

### Upregulation of Ionotropic Receptor genes in the *Orco* mutant legs

We next investigated the potential molecular mechanisms underlying enhanced neuronal activity in *Orco* mutant foreleg neurons. We performed bulk RNAseq from pooled whole forelegs and whole midlegs isolated from wildtype and *Orco* mutant mosquitoes (Figure 7A). Using an FDR cut-off of 0.01, we looked for differentially expressed genes, their expression level, and the degree of differential expression (Figure 7B). We found 614 upregulated and 688 downregulated genes in this *Orco* mutant foreleg and midleg transcriptome dataset.

**Figure 7:**
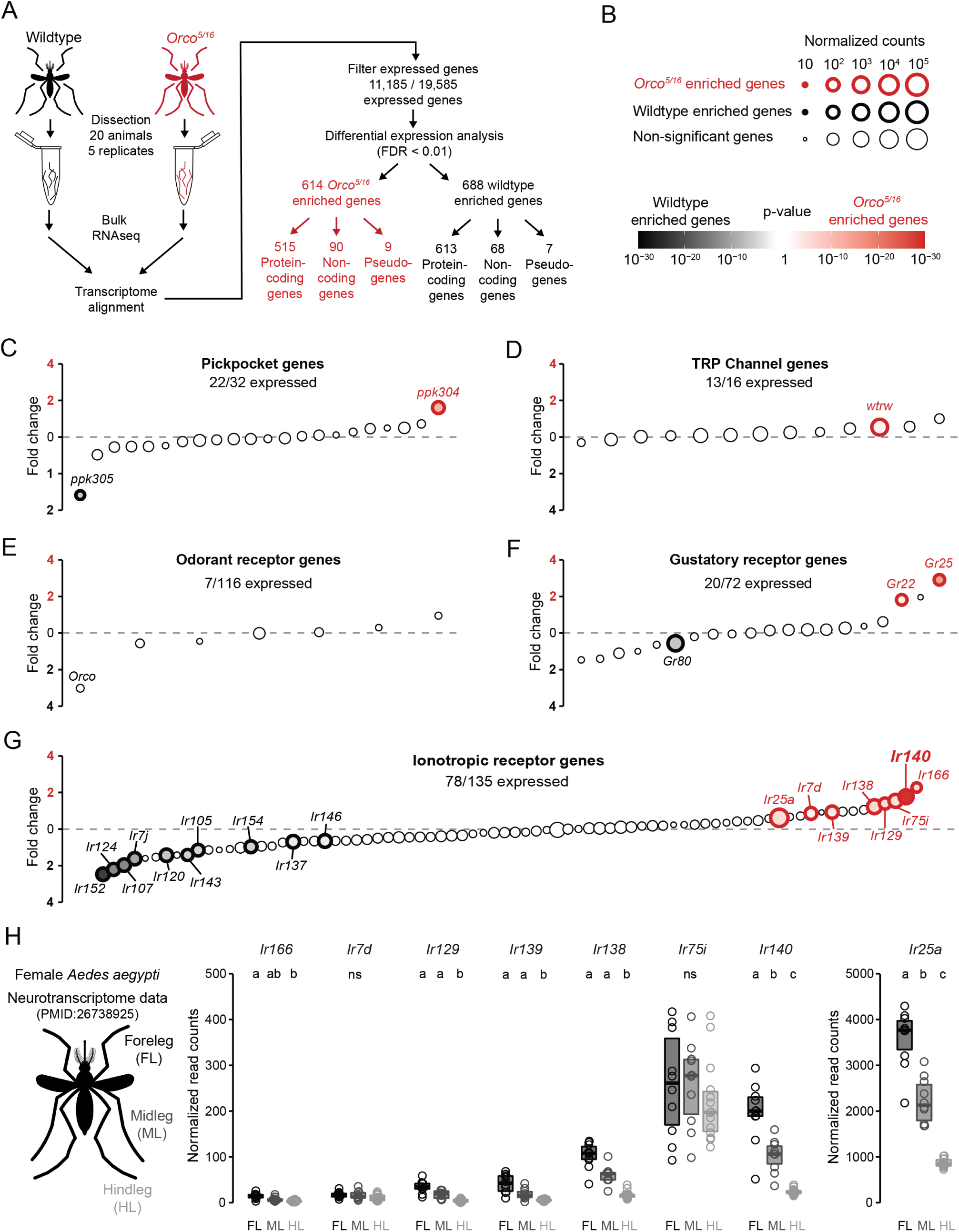
*Aedes aegypti* chemosensory receptor expression changes in the *Orco* mutant legs. (A) Workflow of pooled foreleg and midleg bulk RNA-sequencing comparing gene expression between wildtype and *Orco^5/16^* mutant mosquito tarsi. (B) A legend describing bulk RNA-seq data presented in panels C-G. The number of normalized read counts is presented as size, differentially expressed genes are represented with bold, colored outlines, and the degree of differential expression is represented as a color gradient. Read counts were normalized using DESeq2 median of ratio method. (C-G) Gene expression comparisons between wildtype and *Orco^5/16^* mutant mosquito tarsi for Pickpocket genes (C), TRP channels (D), Odorant Receptors (E), Gustatory Receptors (F), and Ionotropic Receptors (G). Genes are listed from left to right from lowest to highest fold-change enrichment in *Orco^5/16^* tarsi compared to wildtype. (H) *Orco^5/16^* upregulated Ionotropic receptor gene expression in the indicated tissues from the female leg neurotranscriptome dataset (*29*). Data labeled with different letters are significantly different (n = 10-14 replicates/leg, FDR < 0.01, pairwise DESeq2 comparison).

We narrowed our search by focusing on known sensory receptor genes. Twenty-two out of 32 *pickpocket* genes were detected, one upregulated and one downregulated in *Orco* mutant legs (Figure 7C). For TRP channels, 13 of 16 genes were detected, with only *water witch* (*wtrw*) upregulated in *Orco* mutant legs (Figure 7D). Odorant receptors (ORs) were mainly absent in the legs, with only 7 out of 116 detected, and none were differentially expressed in *Orco* mutant legs (Figure 7E). *Orco* transcripts were still detected in our *Orco_5/16_* samples, but at a lower level compared to wildtype controls with the expected 5-bp and 16-bp deletion in the mapped reads at each mutant allele locus, likely reflecting transcripts undergoing nonsense-mediated decay (Supplemental Figure S4A). At the gene-wide comparison level, *Orco* failed to meet our stringent FDR < 0.01 cut-off but still displayed a strong trend towards significant downregulation in the *Orco_5/16_* samples with adjusted P-value at 0.028 and FDR < 0.05 (Supplemental Figure S4B). Only 20 of 72 gustatory receptors (GRs) were expressed, with two upregulated and one downregulated in *Orco* mutant legs (Figure 7G). In contrast, 78 out of 135 IRs were expressed, with eight upregulated, and ten downregulated in *Orco* mutant legs.

These transcriptome data suggest that in the legs, IRs are under stronger regulation due to *Orco* mutation than other chemosensory receptor classes.

To further narrow down the potential genes underlying the enhanced heat-evoked neuronal activity in *Orco* mutant forelegs, we reanalyzed the previously published *Aedes aegypti* neurotranscriptome dataset and quantified which *Orco-*upregulated IR genes were also enriched in the forelegs. From our reanalysis, only *Ir25a* and *Ir140* were enriched in the foreleg (Figure 7H). This enrichment was specific to female samples as the male transcriptome dataset detected both *Ir25a* and *Ir140*, but the foreleg enrichment was not as profound as the female samples (Supplemental Figure S4C). The *Ir25a* expression data are consistent with our *Ir25a* heat-seeking behavioral data, suggesting that *Ir25a* is a core co-receptor subunit required for thermotaxis behavior (Figure 1E).

### *Ir140* mediates heat-seeking behavior and enhanced *Orco* thermosensation

To test the role of *Ir140* in thermosensation, we used an enhanced method of CRISPR-Cas9 genome editing that builds on our original method (*21*) to generate *Ir140* mutant mosquitoes that lacked functional Ir140 ligand-specific IR subunits. We isolated two *Ir140* mutant alleles, *Ir140_144_* and *Ir140_17_*, and used the heat-seeking assay to characterize thermotaxis behavior of heterozygous animals and heteroallelic *Ir140_17/144_* null mutants (Figure 8A, Supplemental Figure S5A-C). *Ir140* mutants displayed a significant reduction in heat-seeking behavior compared to their genetic controls only when the Peltier was set at 36°C (Figure 8B). These data suggest that *Ir140* is a key molecular player in *Aedes aegypti* thermotaxis behavior at human skin temperatures.

**Figure 8:**
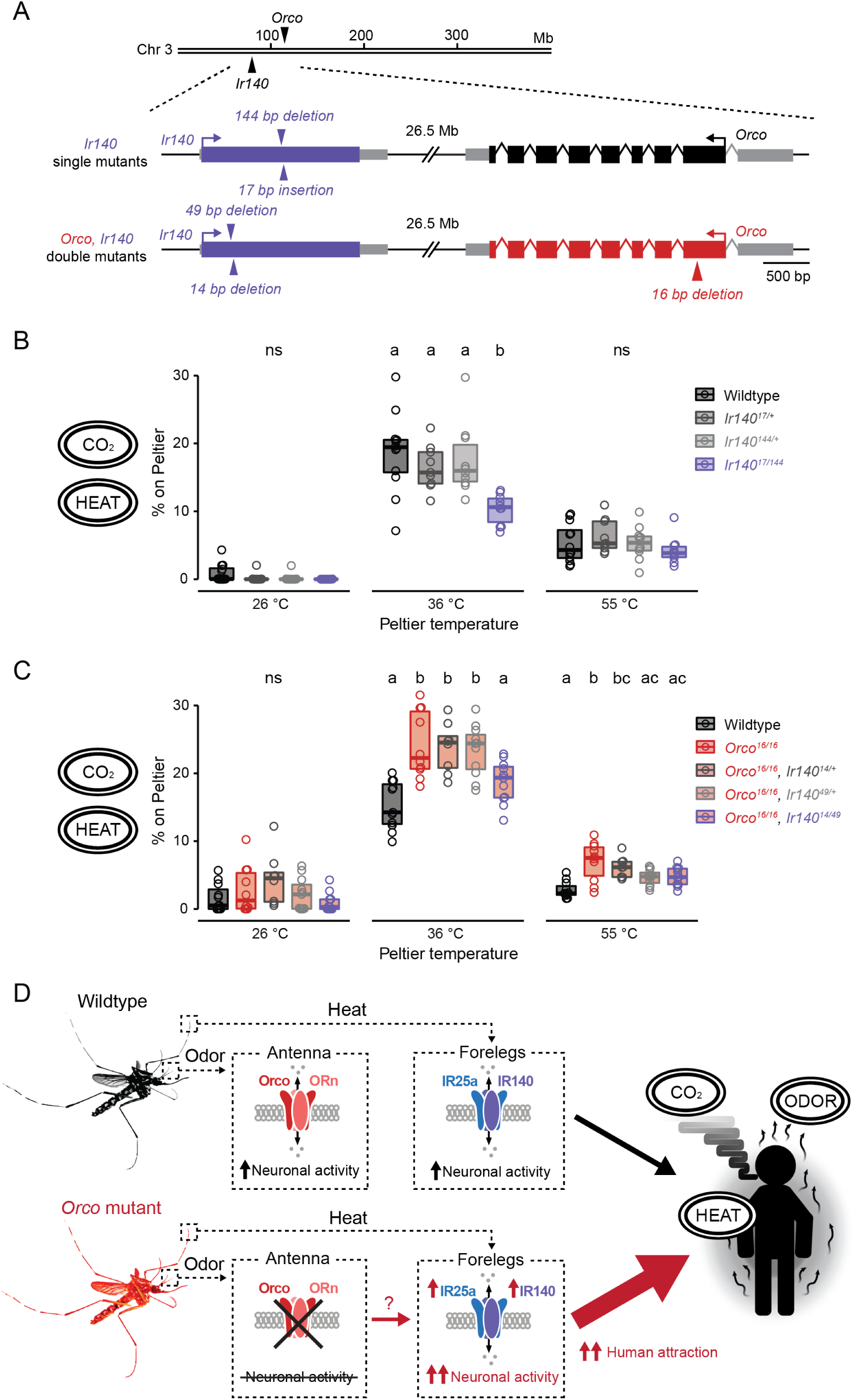
*Ir140* contributes to heat-seeking behavior and is required for the enhanced *Orco* mutant heat-seeking behavior. (A) Schematic of *Aedes aegypti Ir140* and *Orco* loci. Arrows indicate *Ir140* mutant alleles in wildtype and *Orco* mutant backgrounds. Introns are not drawn to scale. (B,C) Percent of mosquitoes of indicated genotype on Peltier during seconds 90-180 of stimuli of indicated temperature for single *Ir140* mutants (B, n = 9-12 trials per genotype) and *Orco*, *Ir140* double mutants (C, n = 8-11 trials per genotype). (D) Summary and model of *Aedes aegypti* heat detection and enhanced thermosensitivity in *Orco* mutant mosquitoes. Data are plotted as scatter-box plots (individual data points, median as horizontal line, interquartile range as box) for (B,C). Data labeled with different letters are significantly different (p < 0.05; one-way ANOVA with Tukey’s HSD post hoc test at each tested temperature).

We further tested the possibility that *Ir140* upregulation could underlie the enhanced thermosensitivity in *Orco* mutants. Because *Orco* and *Ir140* are tightly linked on the 3_rd_ chromosome, it was not feasible to simply cross *Ir140* mutant alleles into the *Orco* mutant background because of the low likelihood of recovering recombination events between the loci. Therefore, we generated *Orco*, *Ir140* double mutants by targeting *Ir140* using CRISPR-Cas9 genome editing in the *Orco_16_* mutant background strain (Figure 8A, Supplemental Figure S5D-F). We isolated two *Orco, Ir140* double mutant alleles, *Orco_16_, Ir140_14_* and *Orco_16_, Ir140_49_,* and tested the thermotaxis behavior of the heteroallelic *Orco_16/16_, Ir140_14/49_* double mutants using the heat-seeking assay. While *Orco_16/16_*, and the heterozygous mutants displayed enhanced heat-seeking behavior, heteroallelic *Orco_16/16_, Ir140_14/49_* double mutant animals failed to show this enhancement and were indistinguishable from wildtype controls (Figure 8C). These results indicate that *Ir140* is one of the key molecular players involved in general thermosensitivity and is required for the enhanced thermosensitivity displayed by *Orco* mutant mosquitoes.

## DISCUSSION

### Cross-modal sensory compensation enhances mosquito attraction to humans

In this study, we establish a novel sensory compensation mechanism in *Aedes aegypti* mosquitoes. Our findings demonstrate that *Orco* mutant mosquitoes exhibit heightened thermosensitivity due to increased neuronal activity in their legs. We further show that many ionotropic receptors were differentially expressed in *Orco* mutants legs. Finally, our study reveals the importance of *Ir140*, a ligand-specific IR subunit, in normal heat-seeking behavior and its necessity for the heightened thermosensitivity observed in the *Orco* mutants (Figure 8D). This cross-modal sensory compensation adds to the repertoire of behavioral strategies *Aedes aegypti* can utilize for their persistent and seemingly unbreakable drive to seek humans. These observations are consistent with prior work showing that disruptions in specific sensory receptors caused reductions in mosquito behavior related to the affected sensory modality but did not eliminate overall attraction to humans (*13, 24, 25, 35*). Co-expression of multiple chemosensory receptor gene families within a single olfactory sensory neuron may permit mosquitoes to rely on at least one chemoreceptor gene family if others become compromised (*26*). It is not known if receptor co-expression is seen in sensory neurons in the leg, the primary thermosensitive organ of the mosquito, but the recent development of techniques for single-cell RNA sequencing of mosquito sensory neurons (*26*) may shed light on this.

### Conservation of temperature detection between *Drosophila* and mosquitoes

Thermosensation has been extensively explored in *Drosophila melanogaster* flies, leading to the identification of numerous genes and sensory organs in detecting cooling and heating temperatures. For noxious heat detection, *painless*, which is expressed throughout the nervous system, is essential for the detection of harmful heat at all life stages (*61, 62*). *Pyrexia* is required for noxious heat-evoked paralysis behavior (*63*). Innocuous warmth detection involves genes including *TrpA1,* required for warmth detection by AC neurons in the brain, controlling the slowly developing preference response of flies exposed to a shallow thermal gradient (*45, 64*). The gustatory receptor *Gr28b(d)* is necessary for rapid noxious heat avoidance (*42, 43*). Detection and avoidance of innocuous cool temperatures relies on *brv1*, *brv2*, and *brv3* expression in cold receptors located in the arista (*44*) and *Ir21a*, along with *Ir93a and Ir25a* in larval dorsal organ cool cells, and in cooling cells located in the adult arista (*65*). *Iav* in the chordotonal organ (*66*) and *Trp* and *Trpl* mutants also display deficits in cool avoidance (*64*). Moreover, *Ir93a* and the IR co-receptor *Ir25a* mediate the detection of warm and cool temperatures (*31, 32*). In addition to ion channels, opsins have been shown to mediate larval temperature preference behavior through *rh5* and *rh6* signaling through the G_q_/PLC pathway coupled to the transduction channel *TrpA1* (*67, 68*).

Recent studies have identified genes conserved from *Drosophila melanogaster* that mediate cooling detection in mosquitoes. In *Anopheles gambiae*, *Ir21a* and *Ir93b* are expressed in sensory neurons at the antennal tip. Mutating these genes significantly reduced neuronal activity in response to cooling air and deficits in heat-seeking behavior at 37°C (*24, 25*). In *Aedes aegypti*, *Ir93a*-expressing neurons in the antenna are thermosensitive, but the role of this ligand-selective IR in heat-seeking behavior remains to be tested (*25*). In contrast to cooling detection, the functional similarity between *Drosophila melanogaster* and mosquito heat detection is not well conserved. Unlike *Drosophila melanogaster TrpA1* mutants, *Aedes aegypti TrpA1* mutants displayed normal thermotaxis behavior in the range of human skin temperature between (26-40°C) but are defective in avoidance of temperatures greater than 40°C, suggesting that *TrpA1* is a noxious heat sensor in mosquitoes (*28*). Additionally, *Gr19*, the *Aedes aegypti* homologue of the *Drosophila melanogaster Gr28b(d)* rapid heat avoidance gene, is not required for thermotaxis behavior (*28*). These findings collectively indicate that while studies in *Drosophila melanogaster* have contributed significantly to our understanding of thermosensation, mechanisms in the fly do not directly translate to the mosquito. This is understandable given the marked difference between the species in how they utilize thermal information. While mosquitoes are attracted to the high temperature of human skin, *Drosophila* strongly avoid these high temperatures. Our current work identified *Ir140* as the first mosquito-specific gene that mediates thermotaxis behavior in *Aedes aegypti*. Based on the complete abolishment of heat-seeking behavior in *Ir25a* mutants, we speculate that additional IR ligand-specific subunits are likely to function as additional heat sensors.

### What is the role of the antenna during thermotaxis behavior?

The mosquito antenna has long been regarded as the essential sensory appendage for mosquito thermotaxis behavior. Earlier studies identified a pair of sensory neurons within the small coeloconic sensilla located at the antennal tip that respond to changes in air temperature (*24, 47–49*). These studies further revealed that one of these sensory neurons responds to rising air temperatures, while the other is sensitive to decreasing temperatures (*47–49*). However, whether the thermosensitive properties of these sensory neurons have any influence over mosquito thermotaxis behavior has never been investigated. An early study demonstrated that removing the antenna in *Aedes aegypti* had minimal impact on their ability to seek a combination of human odor and heat unless more than ten distal antenna segments were removed (*50*). Consistent with these findings, a recent study found that removing the distal six antenna segments in *Aedes albopictus* had little effect on blood-feeding on a human arm (*51*). In this study, we found no evidence that the antenna is required for heat-seeking behavior or that the antenna responds to heat. This is contrary to a recent study showing that *Aedes aegypti* antennal *Ir93a* neurons are thermosensitive (*25*). Although the basis for differences in observations in these two studies is unknown, our data strongly indicate that the thermosensitivity of the antenna tip neurons is not necessary for mosquito thermotaxis behavior. The exact functions of antennal tip thermosensory neurons, if they exist, remains to be determined.

### *Aedes aegypti* tarsi as heat sensors

Insect legs are one of the primary sites for taste detection. In *Drosophila melanogaster*, tarsal sensory neurons detect various non-volatile chemosensory cues, including sugars, bitters, sours salts, fatty acids, water pH, and pheromones (*38, 39, 55, 69*). In *Aedes aegypti*, tarsal neurons play a crucial role in detecting water and salt for oviposition site assessment and in responding to DEET contact repellency (*41, 70, 71*). In this study, we described the first observation of heat-induced activation of tarsal sensory neurons and established a role for the foreleg distal tarsal segments in heat-seeking behavior. On a behavioral level, our findings demonstrate that removing a pair of distal tarsal segments in the forelegs, as opposed to those in the mid-or hindlegs, significantly impairs the mosquito’s ability to detect and land on heated surfaces. At the level of neuronal activity, we show that heat activates a substantial subset of tarsal sensory neurons spanning the entire length of the three most distal segments of both forelegs and midlegs. Furthermore, we present evidence of selective enhancement in heat-evoked neuronal activity within the forelegs of *Orco* mutant animals. These findings indicate the critical role foreleg tarsal neurons play in *Aedes aegypti* thermotaxis behavior.

Removing pairs of tarsal segments from the forelegs significantly reduced heat-seeking and engorgement on the artificial blood feeder, though the behaviors were not completely abolished. We speculate that this residual behavior may be attributed to the redundant thermosensory abilities of the neurons in the midlegs, given that these neurons also exhibited heat-evoked neuronal activity. We could not obtain functional imaging data in the hindleg tarsi due to high background fluorescence in this tissue. Thus, the contribution of hindlegs to heat detection remains to be determined. As established in previous reports, heat dissipates rapidly from its source (*25, 72*). Therefore, mosquitoes must be close to the surface to detect the presence of heat and swiftly assess the surface temperature. The behavioral aspects of mosquito landing events have received limited attention, and future studies using high-resolution behavioral tracking approaches should investigate how each sensory appendage is utilized during heat-seeking behavior.

### IRs regulate mosquito thermosensitivity toward host skin temperature

Our data reveal that functional IR complexes, formed from a unique combination of IR ligand-specific subunits along with co-receptor subunits, contribute to various aspects of heat-seeking behavior. We found that *Ir25a* is required for thermotaxis behavior across all temperature ranges. This suggests that a functional IR complex containing *Ir25a* is crucial for thermosensation. On the other hand, *Ir76b* mutants exhibited heightened sensitivity to milder heat while retaining wildtype responses to higher temperatures. The heightened sensitivity towards temperatures lower than human skin temperature implies that *Ir76b*-containing IR ion channel complexes are involved in adjusting the preferred temperature to a host-relevant range. The mechanism by which such adjustment could be made is unknown, but we speculate that *Ir76b* contributes to a thermosensitive IR complex that is insensitive to lower temperatures, and that its loss releases inhibition on the complex, activating positive thermotaxis.

*Ir8a* mutants display yet another unique heat-seeking behavior, showing enhanced heat-seeking behavior throughout the entire temperature range. This enhancement in *Ir8a* mutants is similar to the heat-seeking behavior seen in *Orco* mutants, except that this enhancement extends across a wider temperature range and is not specific to human skin temperature. *Ir8a*, like *Orco*, is selectively expressed in sensory neurons in head appendages. It will be interesting to learn if *Ir8a* mutants also show selective upregulation of IR subunits in the legs, or whether *Ir8a* mutants utilize another mechanism of sensory compensation. We speculate that *Ir8a* mutants enhance their heat-seeking behavior to compensate for their loss of lactic acid detection, a major chemical component of human sweat odor. Interestingly, *Ir8a* mutants also show enhanced water-seeking behavior while searching for an egg-laying site (*22*). These data suggest that behavioral enhancement extends beyond human-seeking behavior to encompass other crucial tasks necessary for the propagation of their offspring. Future studies are needed to investigate whether the IR signaling pathways tune additional sensory modalities.

### What is the signaling mechanism underlying mosquito sensory compensation?

Sensory compensation has traditionally been associated with localized changes in synaptic connections and strength (*8*), reorganization of sensory maps (*6*), direct neuromodulation at the synapse by opposing sensory modalities (*9*), and the release of neuromodulators from remote brain structures (*10, 11*). However, we believe these mechanisms are unlikely to play a role in the case of *Orco* mutants, given the presence of multiple synapses between the sensory appendages in the head and the affected neurons in the leg. Head appendage sensory neurons synapse in the brain while leg sensory neurons synapse in the ventral nerve cord, the equivalent of the vertebrate spinal cord.

We speculate that this form of sensory compensation arises from a non-cell autonomous mechanism independent of *Orco* expression in the legs. First, Orco-expression is relatively low compared to other classes of chemosensory ion channels. Using normalized read counts from DESeq2 analysis of foreleg and midleg bulk RNA-seq, we found very low levels of the head appendage co-receptors *Orco* (83±40) and *Ir8a* (88±7), compared to the IRs that we speculate play a role in heat-seeking in the leg: *Ir25a* (17,841±500), *Ir76b* (15,621±1023), and *Ir140* (571 ±14). Moreover, we rarely observed Orco-positive cells when using the *Orco-QF2 >QUAS-dTomato* reporter line (data not shown), further supporting the hypothesis that the contribution of Orco to the compensatory mechanism is indirect.

We propose several alternative mechanisms that could mediate the form of sensory compensation in our study. One potential mechanism involves long-range excitability changes due to neuromodulation. Our RNAseq data reveals enrichment of upregulated neuromodulatory receptor gene expression in the legs of *Orco* mutants. These data suggest sustained neuromodulation via circulating hemolymph leading to transcriptional changes and upregulation of thermosensors to enhance sensitivity to heat. Because our functional imaging data involves the examination of detached legs in *Orco* mutants, acute neuronal modulation is unlikely to be happening. It is also possible that modulation occurs through multi-synaptic modulation by descending neurons from the brain to signal the leg deficit of *Orco* via the ventral nerve cord. One future approach to probe the possible role of neural activity in sensory compensation would be to acutely genetically silence all *Orco*-expressing neurons in head appendages and ask if sensory compensation emerges in foreleg sensory neurons.

Another possibility is that these changes occur during development because the *Orco* mutant is a constitutive knockout, leading to a loss of *Orco* across all life stages of the mosquito. The early loss of *Orco* expression and function could have a lasting impact on the neuronal identity and properties of leg sensory neurons. It is worth noting that *Orco* expression has been detected during the early embryonic stage (*73*), and transcriptomic analysis has hinted at a regulatory role for *Orco* expression in ion channel signaling in *Aedes aegypti* embryos (*74*). With the increased sophistication of genetic tools in the mosquito, it may be possible in the future to knock out the *Orco* gene acutely in adult mosquitoes to ask whether sensory compensation occurs, and if it does over what time course after the knock-out of the gene in head appendage neurons.

### Molecular identities of thermosensors in the mosquito legs

In this study, we identified *Ir140* as a critical mediator of heat-seeking behavior, as demonstrated by the significant reduction in heat-seeking behavior observed in *Ir140* mutants. While there is a significant reduction in *Ir140* mutant heat-seeking behavior, the behavior is not entirely abolished. Therefore, we speculate that additional IR subunits may form functional ion channels contributing to heat detection. One of the upregulated ligand-specific IR subunits in *Orco* mutant legs may be among the remaining genes that play a role in mosquito heat-seeking behavior. We speculate that *Ir140*, in conjunction with *Ir25a* and potentially other IR subunits, forms an ion channel within the tarsal sensory neurons. This ion channel complex could serve as either a direct thermosensor or function downstream in the signal transduction pathway of the thermosensory neurons. Further investigation into their biophysical properties using a heterologous expression system is needed to determine whether these ion channel complexes are intrinsically thermosensitive.

### Concluding remarks

In this work, we have discovered a novel mechanism of sensory compensation in the mosquito involving a transcriptional response in the tips of the foreleg in response to the loss of the major olfactory co-receptor in head appendages. We further show that the leg, not the antenna, is the behaviorally relevant appendage for sensing heat during human host seeking. Taken together this work highlights the extraordinary ability of the female mosquito to retain attraction to humans, whose blood is key to the generation of her offspring.

## ACKNOWLEDGMENTS

We thank members of the Vosshall Lab for comments on the manuscript; Libby Mejia, Melissa Dallesandro, and Gloria Gordon for strain maintenance; Román Corfas for technical advice on behavioral assays; Veronica Jové for discussion and advice on functional imaging; Connie Zhao at the Rockefeller Genomics Resource Center; Alison North, Carlos Rico, Christina Pyrgaki, and Tao Tong at the Rockefeller Bio-Imaging Resource Center (RRID:SCR_017791); Mona Liu for early support of mosquito injections as part of optimizing the new CRISPR-Cas9 technique; Madison Walker for mosquito injections to generate the *Ir140* mutants.

This work was supported by a Harvey L. Karp Discovery Award and a Japan Society for Promotion of Science Overseas Research Fellowship (T.M.), Jane Coffin Childs Memorial Fund for Medical Research Postdoctoral Fellowships (T.R.S., B.J.M), Kavli Neural Systems Institute Graduate (O.V.G.) and Postdoctoral (T.R.S.) Fellowship, and is based upon work supported by the National Science Foundation Graduate Research Fellowship under grant No. 1946429 to O.V.G. L.H.Z. is a Junior Fellow of the Simons Society of Fellows; L.B.V. is supported by the Howard Hughes Medical Institute.

## AUTHOR CONTRIBUTIONS

T.M. carried out all experiments in the paper with the following exceptions. N.G.L. conducted experiments and analyzed data in Figure 3B, Figure 4K, and Supplemental Figure S1. T.R.S. carried out experiments and analyzed data in Figure 4K and Supplemental Figure S1. R.K.vH. carried out experiments and analyzed data Figure 5. O.V.G. supervised experiments and analyzed data in Figure 5. M.D. carried out experiments and analyzed data in Figure 2H, I. B.J.M. carried out experiments and analyzed data in Supplemental Figure S2. L.H.Z developed the enhanced CRISPR-Cas9 technique used to develop *Ir140* mutants. T.M. and L.B.V. together conceived the study, designed the figures, and wrote the paper with input from all authors.

## OPEN ACCESS STATEMENT

This article is subject to HHMI’s Open Access to Publications policy. HHMI lab heads have previously granted a nonexclusive CC BY 4.0 license to the public and a sublicensable license to HHMI in their research articles. Pursuant to those licenses, the author-accepted manuscript of this article can be made freely available under a CC BY 4.0 license immediately upon publication.

## Methods

### Human and animal ethics statement

Blood-feeding procedures and mosquito behavior with live human hosts were approved and monitored by The Rockefeller University Institutional Review Board (IRB protocol LVO-0652) and the Rockefeller University Institutional Animal Care and Use Committee (IACUC protocol 20068-H). Human subjects gave their written informed consent to participate in this study.

### Mosquito rearing and maintenance

Mosquito strains were reared at 26°C with 70-80% relative humidity and daily 14-hour light and 10-hour dark cycle (lights on at 7 AM) as previously described (*20*). All behavioral experiments except the thermal animal tracking assay were performed at this temperature. Embryos were hatched in a hatching broth media, which was made by crushing one pellet of fish food (TetraMin Tropical Tablets) using a mortar and that was added to 850 ml deionized water and further autoclaved. The following day, 2500 ml of deionized water was added to the hatched larvae and they were fed two fish food tablets per day until pupation. After eclosion, adult mosquitoes had access to 10% sucrose (w/v in distilled water), which was delivered in a Boston clear round 60 mL glass bottle (Fisher, FB02911944) with the sugar fed through a saturated cotton dental wick (Richmond Dental 201205). Female mosquitoes were blood-fed on mice or human arms for egg collection. Eggs were dried at 26°C and 80% humidity for three days, then stored at ambient temperature and humidity for up to 4 months. All behavior experiments were carried out using adult female mosquitoes that were 7 to 21 days post-eclosion in the light phase of the photoperiod. The *Aedes aegypti* Orlando strain was used as the wildtype control and in the generation of all animal crosses except Figure 1E, G, where the genetic background was the Liverpool IB12 strain.

### Manipulation of sensory appendages

Adult female mosquitoes were immobilized using cold anesthesia and subsequently positioned on a chilled cold block. Sensory appendages were trimmed with 2.5 mm straight-edge Vannas spring scissors. For experiments involving the trimming of antenna tips, the three most distal antenna segments (segments 11-13) were removed from both antennae. When conducting full antenna removal experiments, both antennae were severed right above the Johnston’s organ. For tarsal tip cut experiments, the third, fourth, and the fifth tarsomere were removed from each pair of forelegs, midlegs, and hindlegs. Mosquitoes were allowed to recover overnight at 26°C with 70-80% relative humidity. Mock treated controls underwent cold anesthesia, and the sensory appendages were handled with the Vannas spring scissors, but no trimming was performed.

### Arm-next-to-cage assay

This assay was modified from an arm-next-to-cage assay described previously (*75*). Briefly, for each trial, 20-40 adult female mosquitoes were sorted under cold anesthesia (4°C) and transferred to a 28 cm^3^ cage and allowed to recover overnight at 26°C with 70-80% relative humidity and fasted by replacing 10% sucrose with deionized water. A human forearm was placed adjacent to one side of the cage, separated by two 10 mL serological pipet tips, and perpendicularly to the arm. This arm positioning allowed mosquitoes to detect human odor, CO2, and heat while preventing the mosquitoes from directly making skin contact. A monochrome CMOS camera (The Imaging Source, DMK37BUX178) with a 12 mm/F1.8 lens (Edmond Optics, #33-303) was positioned to take images of mosquitoes responding to the human arm. Trials ran for 5 minutes, and images were acquired at one frame/second using the IC Capture Image Acquisition software (The Imaging Source). We manually counted the number of mosquitoes resting overlaying the human arm to quantify mosquito responses. For the human arm experiment with DEET, 10% DEET (N,N-Diethly-3-methylbenamide; Sigma D100951, mixed with 100% ethanol, v/v) was applied on the entire length of the forearm surface facing the cage. Applied 10% DEET was allowed to dry for 10 minutes before each trial. After drying, behavior experiments were performed as described above.

### Nylon-next-to-cage assay

This assay is a modified version of the arm-next-to-cage assay as above. Instead of a live human arm, a clear 80-well microcentrifuge test tube rack was inserted into a worn or unworn nylon sleeve. Human odor collection using nylon sleeves was described previously (*35*). The nylon sleeves were kept taut using binder clips. This nylon sleeve stimulus was further mounted to a ring stand and placed adjacent to one side of the cage, ensuring that it did not make direct contact with the cage. An eighty-well microcentrifuge test tube rack was used to mimic the surface area of the human arm that mosquitoes encountered during the arm-next-to-cage assay (6.5 cm x 22.5 cm surface). Image acquisition was performed as described in the arm-next-to-cage assay.

### CO_2_ activation assay

Twenty to 40 adult female mosquitoes were sorted under cold anesthesia (4°C) and transferred to a 28 cm^3^ cage and allowed to recover overnight at 26°C with 70-80% relative humidity and fasted by replacing 10% sucrose with deionized water. Sorted mosquitoes were transferred into a custom-made Plexiglass box (30 cm^3^), with carbon-filtered air pumped continuously into the box via a diffusion pad installed on the ceiling of the enclosure. Mosquitoes were allowed to acclimate for 10 minutes. Following acclimation, a 20-second pulse of CO_2_ was applied, and flying activity was monitored for 10 minutes. A monochrome CMOS camera (The Imaging Source, DMK37BUX178) with a 12 mm/F1.8 lens (Edmond Optics, #33-303) was positioned to acquire images at one frame/second using the IC Capture Image Acquisition software (The Imaging Source) of the entire behavior box to capture of mosquitoes flying in response to CO2.

### Heat-seeking assay

Experiments were performed as previously described (*13, 28*). Briefly, 40 – 50 adult female mosquitoes were sorted under cold anesthesia and allowed to recover overnight at 26°C with 70-80% relative humidity and fasted by replacing 10% sucrose with deionized water. Before each trial, sorted and fasted mosquitoes were transferred into a custom-made Plexiglass box (30 cm^3^), with carbon-filtered air pumped continuously into the box via a diffusion pad installed on the enclosure’s ceiling and each stimulus period lasted 3 minutes on a single Peltier element (6 × 9 cm, Tellurex). The surface was covered with 15 × 17 cm standard white letter-size printer paper (NMP1120, Navigator) and held taut by a magnetic frame. CO_2_ pulses (20 seconds, to > 1000 parts per million above background levels) were added to the air stream and accompanied all heat stimulus period onsets. Mosquito landings on the Peltier were monitored by fixed cameras (FFMV-03M2M-CS, Point Grey Research), with images acquired at one frame/second. Images were analyzed using custom MATLAB scripts to count mosquito landings within a fixed target region. Mosquito occupancy on the Peltier was quantified during seconds 90–180 of each stimulus period.

### Heat-seeking assay with DEET

To test if DEET could disrupt mosquito attraction without host odor, we tested the female mosquito’s thermotaxis to a 40°C Peltier element in the presence or absence of DEET. A standard 28 x 28 x 28 cm rearing cage (Bioquip) was placed inside a clear vinyl bag. The bag and cage were positioned adjacent to a PCR machine so that the Peltier element was approximately 1 cm from the cage screen. A Firefly MV camera was positioned to take stereotyped images of mosquitoes responding to the heat source (Point Grey Research). A Flypad (Flystuff.com) was placed inside the bag, on top of the cage, as a source of carbon-filtered air and CO2. The assay was assembled on a metal peg board to maintain the fixed position of the components: a rearing cage inside a vinyl bag, a camera, and a PCR machine. Ten to thirty minutes prior to the start of the assay, 25 adult female mosquitoes were released into each cage to acclimate. Before starting each trial, 100 µl of 100% ethanol or 10% DEET in ethanol was applied to a 2 cm x 6 cm strip of Whatman filter paper (GE Healthcare). The treated paper was hung from a rectangular plastic frame in front of the Peltier element, so it was not in contact with the cage screen. For each trial, carbon-filtered air was pumped into the assay for 60 minutes to remove the host odor. After the 60-minute pre-stimulus period, 10% CO_2_ was released using the Flypad while the Peltier element was heated to 40°C. For each 5-minute trial, images were taken every 10 seconds. After each trial, the mosquitoes were placed under cold anesthesia and counted to verify the number used. To quantify mosquito behavioral responses in this assay, we counted the number of mosquitoes on the cage screen near the Peltier element. Each data point represents a percentage of mosquitoes attracted to the heat source. This was determined by the average number of mosquitoes from the last ten images of each trial divided by the total number of mosquitoes in the assay, multiplied by 100. All scored images were cropped to the same size with Fiji software using a macro [makeRectangle(269, 176, 347, 237); run(“Crop”); run(“Save”); run(“Open Next”)]. Once cropped, images were visually scored with assistance from the multipoint counting tool in Fiji.

### SciTracks assay

The assay was performed as previously described (*13*). A multi-insect 3D tracking system was custom-designed and built in collaboration with SciTrackS GmbH (Sci-Trak). It consists of a flight chamber (1.25 m wide x 1m high x .75 cm deep) with clear acrylic sides and front. A pump continuously provides carbon-filtered air (Cole-Parmer Quiet Pressure Pump #79610-81). CO_2_ can be added to this airstream through the use of a computer-controlled solenoid valve, a valve controller (NResearch, Inc. model 360D1X75R) operated by a MATLAB script through an analog/digital interface (Measurement Computing Corporation, model USB-1208FS). Humidity was controlled using a closed-loop humidification system consisting of an ultrasonic humidifier (SPT model SU-1051B) controlled by a temperature and humidity probe attached to a microcontroller. Humidity remained at 45% +/-4% for the duration of each trial. Tracking is accomplished by imaging the chamber with two offset cameras (Basler piA640-210gm; Basler AG, Ahrenburg, Germany with 1/2 4-12mm F/1.2 IR Aspherical objects, Tamron, Commack, NY, USA). Lighting is provided by two banks of IR LEDs (Metaphase model ISO-23-IRN-24-AL2, Metaphase Technologies, Bensalem, PA). 3D position information of each visible mosquito was calculated in real-time at 200Hz using custom-built software based on previously published algorithms (*76*). Tracks were further processed, filtered, and analyzed using MATLAB. CO_2_ activation: For each trial, 20 female mosquitoes (5-10 days old, fasted overnight with access to water) were added to the flight chamber with a mechanical aspirator. After acclimation and tracking baseline activity for 30 minutes, a 40 second pulse of 100% CO_2_ was added to the airstream at a flow rate of 1360 mL/minute. This raised the concentration of CO_2_ to approximately 1%, as measured in the middle of the flight chamber. (CARBOCAP® Hand-Held Carbon Dioxide Meter GM70, Vaisala Inc., Woburn, MA, USA). Tracking continued for another 20 minutes to measure activity in response to this pulse of CO2. Data are presented as the population distance tracked in 10 second bins or as cumulative distance tracked per mosquito in the 6 minutes immediately before or after CO_2_ addition.

### Optothermocycler assay

The Optothermocycler assay was performed as previously described (*53*). The assay was constructed on top of a PCR thermocycler (Eppendorf Mastercycler) using optomechanical components (Thorlabs). Green light at a wavelength of 530 nm was emitted using six Luxeon Star SP-01-G4 LEDs under the control of an Arduino Uno board. The surface of the PCR block was covered with black tape to minimize glare (Thorlabs T137-2.0). Temperature monitoring was conducted through a type T thermocouple (Harold G Schaevitz Industries LLC CPTC-120-X-N), which was connected to an Arduino board (Arduino A000066) via a thermocouple amplifier (Adafruit MAX31856). The thermocouple sensor was securely positioned on the lower right surface of the PCR block using black tape (Thorlabs T137-2.0). Using a custom Processing script, temperature measurements, and light output were logged at 100-millisecond intervals through the Arduino. The video was synchronized with the light and temperature stimuli by employing an infrared 940 nm LED (Adafruit 387) positioned within the camera’s field of view. Mosquitoes were illuminated with an 850 nm infrared LED strip (Waveform Lighting) surrounding the mosquitoes’ plate orthogonal to the camera’s view. The video was recorded using a Blackfly camera (FLIR BFS-U3-16S2M-CS) outfitted with a 780 nm long pass filter (Vision Light Tech LP780-25.5) at 30 frames/second using Spinview software (Teledyne FLIR). The PCR thermocycler was programmed to apply heat stimuli directly from ambient to a noxious temperature range of 24-60°C.

### Arm feeding assay

Female adult mosquitoes were cold-anesthetized, and 10 – 20 mosquitoes were sorted into a behavioral container made from a 32-ounce HDPE plastic cup (VWR #89009-668). The behavioral container was prepared by cutting a 10 cm hole in the lid with a razor blade and covering the cup with a 20 cm x 20 cm piece of white 0.8 mm polyester mosquito netting (American Home & Habit Inc. #F03A-PONO-MOSQ-M008-ZS) and securing the mesh to the cup using the modified lid. Sorted mosquitoes were allowed to recover overnight at 26°C with 70-80% relative humidity and fasted by replacing 10% sucrose with a cotton wick saturated with deionized water placed on top of the container across the mesh netting. For each experimental trial, mosquitoes were allowed to acclimate for 10 minutes, and a live human arm was placed on top of the container. Mosquitoes were allowed to feed on the human arm through the mesh netting for 10 minutes and were then anesthetized at 4°C and scored as fed if any level of feeding was observed, as assessed by visual inspection of the abdomen of the animal.

### Glytube feeding assay

Female adult mosquitoes were sorted into individual behavioral containers as described with the arm feeding assay. The assay chamber was a modification of previously published methods (*58*) and used a translucent polypropylene storage box 36 cm L x 31 cm W x 32 cm H with a removable lid. One 1.5 cm hole was made on the chamber wall and was used to introduce silicone tubing for CO_2_ delivery. The CO_2_ diffusion pad (8.9 cm x 12.7 cm; Tritech Research) was affixed to the inner center of the lid to allow delivery of purified air and CO_2_ to condition the chamber atmosphere during the trial. Females were fed sheep blood (Hemostat DSB100) supplemented with ATP (Sigma A6419) at a final concentration of 1 mM using a Glytube membrane feeder (*58*). All blood meals and Glytubes were preheated for at least 10 minutes in a 45°C water bath, and ATP was added to meals immediately before feeding. At the start of each trial, up to four cups were placed in the assay chamber, and the animals were allowed to acclimate for 10 minutes before 1 Glytube containing 1.5 mL of sheep blood+ATP mixture was placed on top of each cup, and CO_2_ was turned on for 10 minutes. Mosquitoes were then anesthetized at 4°C and scored as fed if any level of feeding was observed, as assessed by visual inspection of the abdomen of the animal.

### RNA extraction and sequencing

Seven to 10-day-old female wildtype and *Orco^5/16^* mosquitoes were cold-anesthetized and kept on ice for up to 30 minutes or until dissections were complete. Forelegs and midlegs were dissected and collected on ice in a 1.5 mL tube. Five samples from each genotype were collected, and each replicate consisted of legs from 20 animals.

After collecting each sample, tubes were immediately snap-frozen using a cold block (Simport S700-14) chilled with ethanol containing dry ice. Dissected tissue was stored at −80°C until tissue collection was complete. RNA was extracted by homogenizing each sample in TRIzol reagent (Invitrogen 15596026) using a handheld homogenizer (DWK Life Sciences 749540-0000). To the homogenized sample, 20% of the total TRIzol regent volume of chloroform:isoamyl alcohol (Fisher Scientific AC327155000) was added for phase separation. The aqueous phase containing total RNA was transferred into a fresh RNAse-free tube and cleaned using the PicoPure Kit (ThermoFisher KIT0204) following the manufacturer’s instructions, including the DNase treatment step. Samples were run on a Bioanalyzer RNA Pico Chip (Agilent 5067–1513) to determine RNA quantity and quality. Libraries were prepared using the Illumina TruSeq Stranded mRNA kit (Illumina 20020594). Samples were pooled, and sequencing was performed at The Rockefeller University Genomics Resource Center on an Illumina NovaSeq 6000 sequencer using V1.5 reagents, the SP flow cell, and NovaSeq Control Software V1.7.0 to generate 150 bp paired-end reads. All samples were sequenced on the same flow cell to correct for potential batch effects. Data were demultiplexed and delivered as fastq files for each library. Sequencing reads have been deposited at the National Center for Biotechnology Information (NCBI) Sequence Read Archive (SRA) under BioProject PRJNA1020561.

### Transcriptome alignment, quantification, and differential gene expression analysis

The sequencing data was aligned and quantified with the Salmon quantification software (*77*) using the reference transcriptome generated from *Aedes aegypti* LVP_AGWG (https://vectorbase.org/vectorbase/app/record/dataset/DS_cc8d875d2e) genome and the updated version of the previously published gene annotation file from Jove et al. (*58*). The most current version of the reference transcriptome is available at https://github.com/VosshallLab/Morita_Vosshall2023. Gene expression levels as raw read counts and abundances were retrieved using tximport (version 1.22.0) (*78*). Normalization of raw read counts and differential expression analysis was performed using DESeq2 (version 1.34.0) (*79*). The false discovery rate was calculated using Benjamini & Hochberg multiple testing correction, using the FDR cut-off at 0.01. Foreleg, midleg, and hindleg samples from the neurotranscriptome dataset were reanalyzed following the above steps (*29*). Leg samples from sugar-fed and ovipositing samples were used for analysis (SRR1167521-533, SRR1166473-484, SRR1167469-477, SRR1167554-562, SRR1167491-492). Pair-wise differential expression analysis using DESeq2 was performed. The false discovery rate was calculated using Benjamini & Hochberg multiple testing correction, using the FDR cut-off at 0.01.

### Genome alignment and visualization

The sequencing reads were first processed using Trim Galore (version 0.6.7) to apply library adapter and quality trimming (https://zenodo.org/record/8381219). Trimmed reads were aligned to the *Aedes aegypti* LVP_AGWG genome using STAR aligner (version 2.7.6a) (*80*). Aligned reads were visualized using IGV (*81*).

### Visualization of reporter line-expressing cells in tarsus and cell counting

We visualized sensory neurons in the fifth tarsomere of the tarsus as previously described (*71*). We used the following reporter lines *brp-T2A-QF2w* (*57*), *Ir25a-T2A-QF2* (*26*), *Ir76b-T2A-QF2* (*26*) crossed to *15xQUAS-dTomato-T2A-GCaMP6*s (line #5) animals (*71*). *Gr4-T2A-QF2* (*58*) was crossed to *15xQUAS-dTomato-T2A-GCaMP6*s animals (line #1) (*58*). There was no difference in the number of cells labeled with dTomato when either *15xQUAS-dTomato-T2A-GCaMP6*s (line #5) or *15xQUAS-dTomato-T2A-GCaMP6*s (line #1) animals were crossed to *brp-T2A-QF2w* (data not shown). Animals were anesthetized on wet ice. Entire tarsi were removed and placed in ice-cold ethanol for approximately 20 seconds and then mounted onto a prepared slide with 50 μL of glycerol pipetted using a 200 μL pipette tip with the end cut to prevent bubbles. A 1.7 mm coverslip was mounted, and pressure was gently applied before sealing with nail polish. Confocal z-stacks were acquired on a Zeiss Inverted LSM 880 laser scanning confocal with a 40x/0.8 NA water immersion corrected objective at a 1024 x 1024 pixel resolution and a voxel size of 0.2076 μm x 0.2076 μm x 1 μm. Cranial and caudal neuron layers were acquired in the order of proximity to the objective, which was random based on the leg’s orientation. Cranial and caudal layers were identified based on whether the leg imaged was the left or right leg, and relative to the position of the claw. The cranial and caudal layers are easily distinguished by a space between the cranial and caudal layers that rarely, if ever, contains labeled neurons and also includes the trachea. Whichever layer was further away from the objective was typically dimmer. Laser intensity correction was applied to ensure the distal layer received enough laser power, and the proximally imaged layer was not saturated. Cells were counted using Imaris software (Bitplane). The 3D image was cropped for each image to include the entire fifth tarsomere. Cells were counted first using an automated counting algorithm, then each cell was confirmed manually. The estimated spot diameter for the automated counter was 5 μm. All cells were counted for each collected sample. For distinguishing cell counts in cranial and caudal layers on an individual sample or counting for 50 μm distal-proximal segments of the fifth tarsomere, clipping planes were used on the 3D image so that no individual cells in a given sample were counted twice. Fluorescent sensilla counting was done manually. The side view of the cranial and caudal layers of the tarsus was achieved by rotating a 3D image of a confocal Z-stack.

### *Ex vivo* sensory appendage imaging

Calcium imaging was performed on a microscope setup previously described (*58*). Experiments were performed on an inverted Ti-2E wide-field microscope (Nikon) with a dual FITC/TRITC bandpass cube and alternating emission wheel with 520/40 GFP and 628/40 RFP bandpass filters. An nd2 filter was added with the 628/40 RFP bandpass filter to attenuate the dTomato signal. Images were acquired with a 20x/0.75 air objective (Nikon) and Zyla 4.2 Plus camera. Calcium imaging experiments were performed on female mosquitoes 7–14 days post-eclosion. The imaging chamber was prepared by affixing a Gold Seal Cover Glass, No. 1 22 × 40 mm coverslip (Ted Pella #260353) to a low profile open diamond bath imaging chamber (Warner Instruments RC-26GLP) using a silicone lubricant (Dow Molykote 111 O-Ring Silicone Lubricant). Female mosquitoes were anesthetized briefly at 4°C for dissection. Each dissected sensory appendage was removed using fine forceps and Vannas spring scissors. Dissected sensory appendages were placed on a 15 mm round cover glass (Warner Instruments, CS-15R) using a very thin layer of silicone lubricant. This sensory appendage preparation, along with the coverslip, was directly secured inside the imaging chamber. dTomato fluorescence was examined before each trial to locate and focus on the sensory appendage intended for imaging. During the experiment, each image acquisition step captured one GCaMP image and one dTomato image separated by less than the 100 milliseconds required to switch the emission filter wheel. Image acquisition was triggered at approximately one frame per second for each channel.

### Temperature delivery and monitoring

Temperature stimulus was delivered to the dissected sensory appendage using a continuous flow of carbon-filtered air supplied by an air pump (Welch, 5078S). The air was passed through two rounds of deionized water using a gas washing bottle (Chemglass CG111206) to filter and moisten the air stimulus. This main air stimulus was further bifurcated into two independent air lines, and the opening/closing of each opening was controlled using its dedicated solenoid valves (Warner Instruments, VCS-8-PTFE), one for ambient air and another for heated air. An ambient temperature air line outlet was placed adjacent to the dissected sensory appendages. This ambient air line functioned as the default air line and was kept open at the beginning and the end of each experiment. As for the heated air line, tubing was connected to a dual in-line solution Peltier heater/cooler device (Warner Instruments, SC-20), and the outlet was placed directly above the sensory appendage to be imaged. The air temperature was adjusted by controlling the Peltier heater/cooler device temperature using an external bipolar temperature controller (Warner Instruments, CL-100). By default, this heated air line is kept closed unless a heated stimulus is applied to the sensory appendages. The experiment started with the ambient air line opened, and the heat air line closed. The ambient air line was closed for each heat stimulus while the heat air line was opened. When the temperature of the sample reached ∼33°C, the ambient air line was opened while the heat air line was closed. The opening of each air line was controlled through analog signals provided by the Image acquisition software. For the slow heat ramp protocol used in Figure 6H-N, the temperature stimulus was delivered to the dissected sensory appendage through a single air line with an air source as previously described. This air line was passed directly through the Peltier device, where the temperature can be changed throughout the experiment. At the beginning of each trial, the Peltier device was set at 20°C for ambient air temperature delivery. During the experimental trial, the Peltier temperature was changed to 50°C, so the target temperature that the sensory appendages were experiencing was around 33°C. Once the temperature reached our target temperature, the Peltier temperature was switched back to 20°C. To monitor the temperature of the sensory appendages, a thermocouple microprobe was placed adjacent to the samples (Physitemp, IT-23) and was recorded using a digital thermometer (Physitemp, BAT-12) during the entire experiment.

### Calcium imaging data analysis

Fluorescent intensities emitted by fluorescent proteins are temperature-dependent and have an inverse relationship with temperature (*82, 83*). To account for this biophysical property of fluorescent proteins, we took advantage of animals co-expressing GCaMP6s and dTomato in the same cell by capturing images at both fluorescent channels. dTomato functioned as a reference imaging channel to correct for both movement artifacts and temperature-induced changes in fluorescence intensity. Regions of interest (ROI) were marked using the dTomato signal captured at the beginning of each experiment. ROI information was overlaid with all image frames captured during each experimental trial, and fluorescent intensities were recorded for each time point from both GCaMP and dTomato imaging channels. The ratio of GCaMP and dTomato was used to report the temperature-dependent changes in fluorescent intensities at each time point and was normalized to the first frame to report the data as ΔR/R.

### *Ir140* crRNA design

*Ir140* single and *Orco, Ir140* double mutants were generated using methods described previously (*21*). sgRNA sequences were designed with the CHOPCHOP sgRNA design tool (https://chopchop.cbu.uib.no/) (*84*) using the following parameters: In, Aedes aegypti (AaegL5.0); Using, CRISPR/Cas9; For, knock-out, with Ir140 CDS sequence (NM_001358695.1) as the target sequence. For each type of *Ir140* mutants, pairs of target sequences were selected for targeted double-stranded break-induced mutagenesis, with each pair flanking roughly 250 base pairs. The following sequences were used for each type of *Ir140* mutant generation:

*Ir140* single mutant:

1. CAGAAGTTATCATAGCACCTGTTTTAGAGCTATGCT
2. TCTCCATCATGCCAACACACGTTTTAGAGCTATGCT

*Orco^16^, Ir140* double mutant:

1. CCGCCTTCATAATCAATCAGGTTTTAGAGCTATGCT
2. CTCATCGTGATCTACTCTGAGTTTTAGAGCTATGCT

*Orco^5^, Ir140* double mutant:

1. CCGCCTTCATAATCAATCAGGTTTTAGAGCTATGCT
2. CTCATCGTGATCTACTCTGAGTTTTAGAGCTATGCT

### CRISPR injection mix

2 µl of 100 µM Alt-R CRISPR-Cas9 tracrRNA (IDT. #1072532) was mixed with 1 µl of each custom-designed Alt-R CRISPR-Cas9 crRNA (100 µM) and 1 µl of Nuclease-Free Duplex Buffer (IDT. #11-01-03-01). This initial mix was then incubated at 95°C on a thermocycler for 5 minutes, followed by 5 minutes at room temperature. After both incubations, 3µl of Alt-R™ S.p. Cas9 V3, glycerol-free, at 1µg/µl (IDT #10007806) and 2 µl of nuclease-free water were added. The solution was incubated at room temperature for 5 minutes and then placed on ice until the time of injection.

### Injection procedure

Mosquito embryos were injected in-house using a Sutter Instrument injection system (Product # FG-BRE, FG-MPC385) and a Zeiss SteREO Discovery stereo microscope (#495007-9880-010). Embryo injection was carried out as previously described (*85*). About 100 eggs were injected per injection session 60 minutes after laying them. A pre-pulled (Sutter Instrument, #FG-P1000) and pre-beveled (Sutter Instrument, #FG-BV10-D) aluminosilicate glass with Filament (Sutter Instrument. #AF100-64-10) was loaded with the injection mix using a Microloader™ tip (Eppendorf. Cat # 930001007). The eggs were injected on their posterior side and visually examined for slight expansion during injection to endure the transfer of the injection solution. In total, approximately 505 embryos from wildtype *Aedes aegypti* Orlando strain were injected for the generation of *Ir140* single mutant strains, while 490 *Orco^5/5^* and 510 *Orco^16/16^* embryos were injected for the generation of *Orco, Ir140* double mutants. After each injection session, the eggs were recovered from the slide and placed on a wet filter paper in a cup with wet cotton on the bottom to maintain humidity during recovery. The cup was covered with tissue paper and kept for three days in the insectary (27°C, 80% relative humidity) until the eggs were transferred to a hatching solution.

### Screening the injected lines

The hatching rate following injection was approximately 2-20%. G0 females from each injection were crossed with males from their respective background strains, and their G1 offspring were screened for germline mutation by PCR amplification and Sanger sequencing of the regions flanking the sgRNA target sites. Two unique stable mutant lines, each resulting in an early stop codon due to a frameshift mutation, were isolated from each injection except for the *Orco^5^* injection, which had a low hatching rate, and we did not find any isolates with signs of gene modification. For the *Ir140* single mutant, one isolated mutant line had a 144-bp out-of-frame mutation (*Ir140^144^*), and the other had a 17-bp insertion (*Ir140^17^*). For *Orco^16^, Ir140* double mutant, one isolated mutant line had a 49-bp deletion (*Ir140^49^*), and the other had a 14-bp deletion (*Ir140^14^*). Virgin females from each mutant line were backcrossed to their respective genetic background males for three generations before establishing stable homozygous lines. Homozygous mutant lines were intercrossed to generate heteroallelic mutants tested in all behavior experiments alongside appropriate genetic controls.

### *Ir140* mutant genotyping

Genotypes were confirmed using Phire Tissue Direct PCR Master Mix (Thermo Fisher, F170L) followed by gel electrophoresis and Sanger sequencing (Genewiz) using the following primer combination for each mutant allele: *Ir140^144^* and *Ir140^17^*single mutants

Forward: ACGCCACACCCGTTCAAATC Reverse: CGATTTGACTAGCTCCGGAATTG

*Orco^16^, Ir140^49^* double mutant

Forward: TGCTAAGCCACATCAACAAG Reverse: ACGTAGTTGAAGCGCCATTTG

*Orco^16^, Ir140^14^* double mutant

Forward: AGCCGGTCGTCAATCATTAC Reverse: ACGTAGTTGAAGCGCCATTTG

The *Ir140^144^* single mutant allele was detected by a 144-bp deletion using standard electrophoresis protocol (Wildtype band: 515 bp; *Ir140^144^* band: 317 bp) while *Orco^16^, Ir140^49^* double mutant allele was detected by a 49-bp deletion on a 2% agarose gel with a low-voltage (90V) electrophoresis protocol (Wildtype band: 250 bp; *Orco^16^,* Ir140^49^ band: 210 bp). For mutants with small deletions, the presence or absence of endogenous restriction enzyme target sites was used to distinguish between mutant and wild-type alleles. PCR products were generated and digested with the indicated enzyme, producing the indicated bands in mutant and wild type:

*Ir140^17^* single mutant with *StyI*

Wildtype: 329 and 186 bp; *Ir140^17^*: 532 bp *Orco^16^,Ir140^14^* double mutant with *AvaII* Wildtype: 151 and 99 bp;

*Orco^16^,Ir140^14^*: 236 bp

*Orco^16^* mutant allele for *Orco^16^,Ir140^14^* and *Orco^16^,Ir140^49^* double mutants were independently validated using the following primers:

Forward: CCGCACGCTGGGCATCTGGAATC

Reverse: ACGGATAGCACTGTAGTCACCAT

All genotyping experiments were performed with a no DNA control and fragment size validation using a 1Kb Plus DNA ladder (ThermoFisher Scientific, 10787026). See Supplemental Figure S5.

### Quantification and statistical analysis

All statistical analyses were performed using R version 4.1.3 (*86*). Details of graphical representations and statistical methods are reported in the figure legends.

### Supplemental data and figures

Five Supplemental figures and four videos accompany the paper.

### Data and Code availability

Data and analysis code used in this publication is available at GitHub https://github.com/VosshallLab/Morita_Vosshall2023 and Zenodo https://zenodo.org/record/8381219. Sequencing reads have been deposited at the NCBI Sequence Read Archive (SRA) under BioProject BioProject PRJNA1020561 https://www.ncbi.nlm.nih.gov/bioproject?term=PRJNA1020561&cmd=DetailsSearch&log$=activity.

## SUPPLEMENTAL FIGURES

**Supplemental Figure S1.**
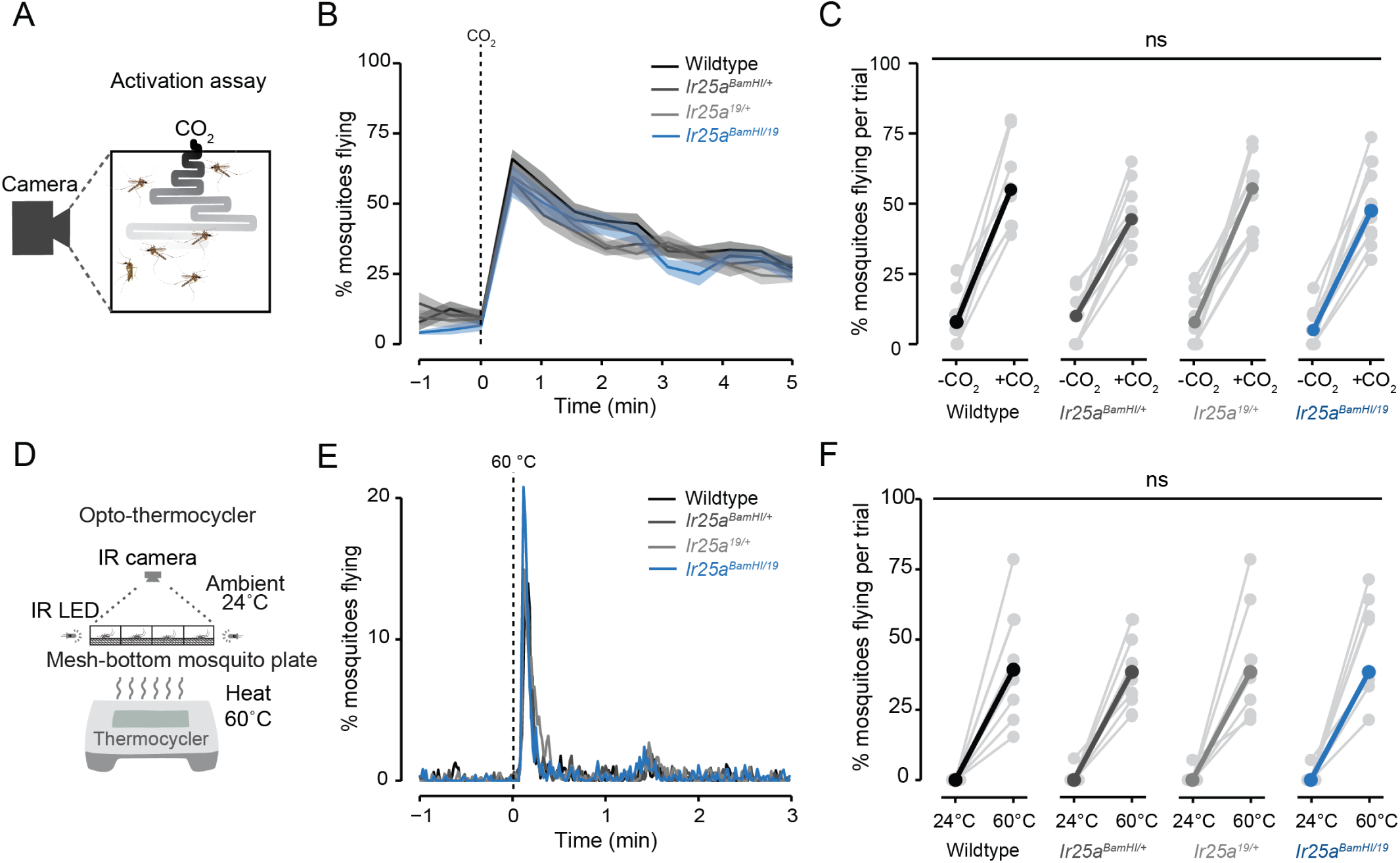
Related to Figure 1 *Ir25a* mutants display normal CO_2_ and noxious heat detection. (A) Schematic of the CO_2_ activation assay. (B) Percent of mosquitoes flying in the CO_2_ activation assay for the indicated genotypes (n = 8-9 trials/genotype. (p > 0.05, between all genotypes tested, one-way ANOVA with Tukey’s HSD post hoc test). Data are presented as mean ± SEM. (C) Quantification of mosquitoes flying during each experimental trial before and after 1 minute after a 20-second pulse of CO_2_ for the indicated genotypes (n = 8-9 trials/genotype, p > 0.05, between all genotypes post CO2, one-way ANOVA with Tukey’s HSD post hoc test). (D) Schematic of the Opto-thermocycler assay. (E) Percent of mosquitoes flying in response to 60°C heat for the indicated genotypes (n = 7-8 trials/genotype). (F) Quantification of mosquitoes flying during each experimental trial before and after 1 minute after a 4-second pulse of 60°C heat for the indicated genotypes (n = 7-8 trials/genotype, p > 0.05, between all genotypes, post 60°C heat, one-way ANOVA with Tukey’s HSD post hoc test).

**Supplemental Figure S2.**
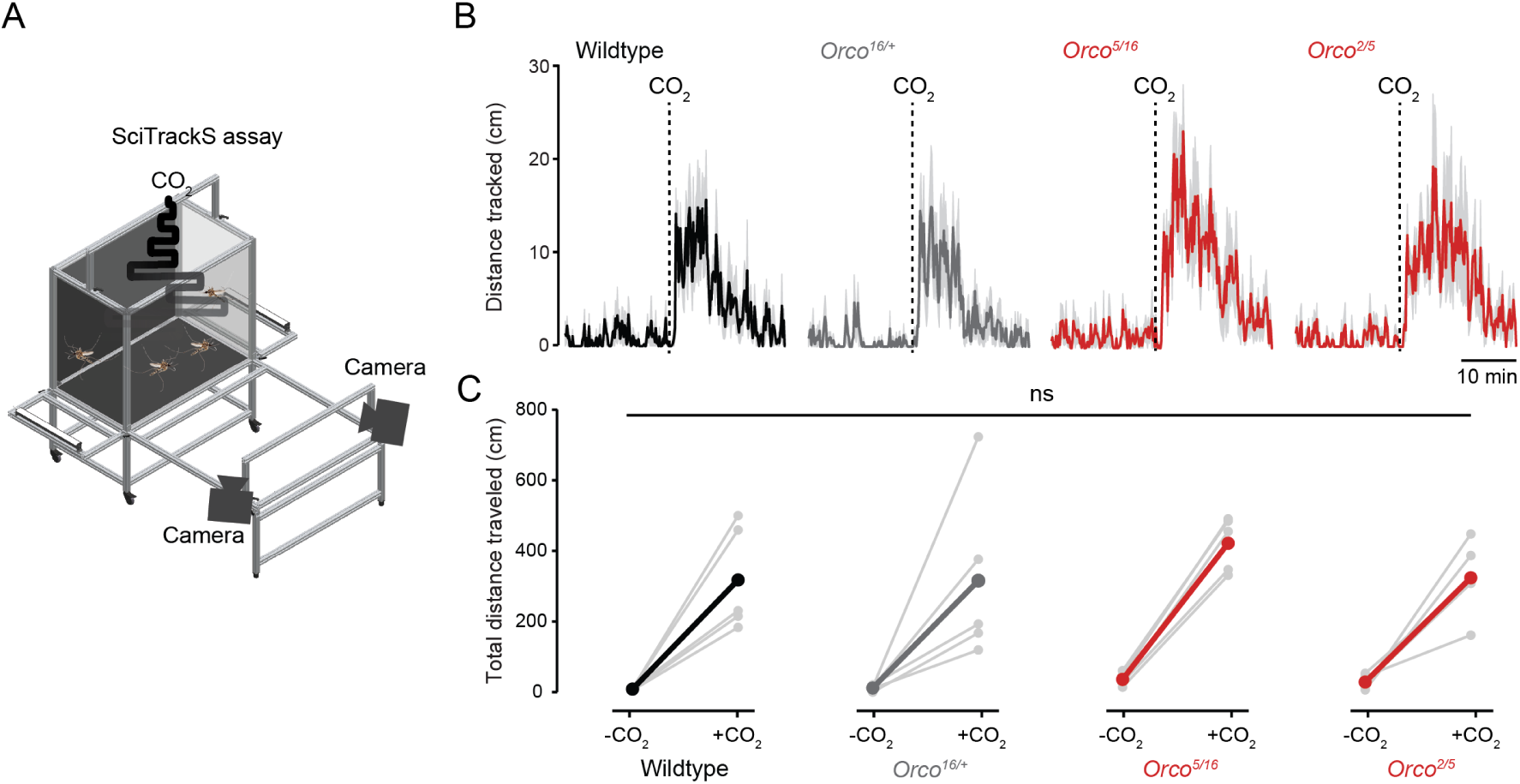
Related to Figure 3 *Orco* mutants display normal CO2-evoked activity. (A) Schematic of the SciTrackS assay. (B) Population distance flown in response to stimulation with CO_2_ for the indicated genotypes (mean ± SEM, 10-second bins, n = 5 trials, 20 animals/trial). (C) Cumulative distance tracked per mosquito in the 6 minutes immediately before or after CO_2_ addition (n = 5 trials/genotype).

**Supplemental Figure S3.**
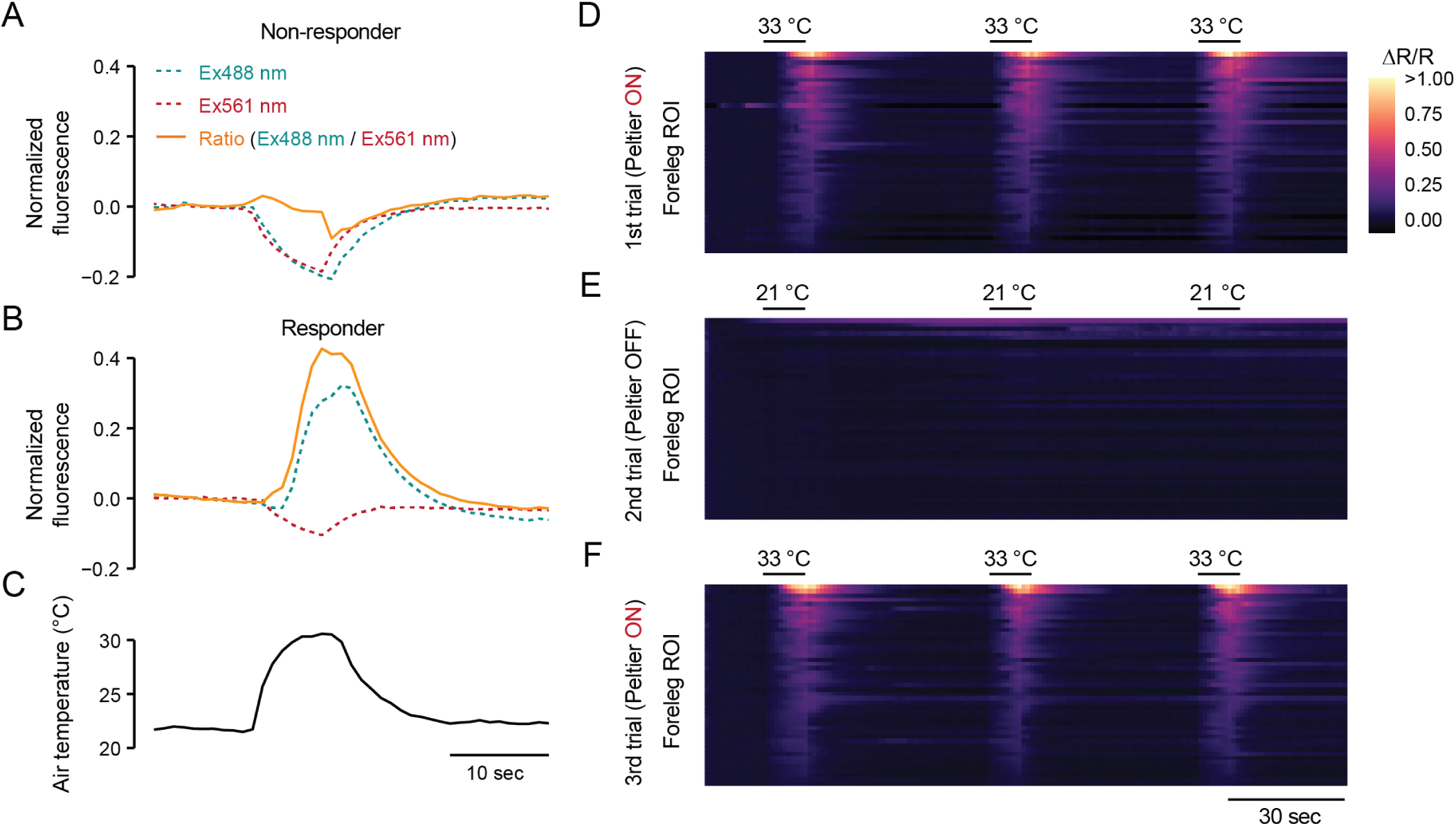
Related to Figure 6 Ratiometric Ca^2+^ imaging analysis. (A-B) Representative fluorescent traces from green, red, and ratiometric channels were acquired from a non-responding (A) and a responding cell (B) in response to a heat pulse (C). (D-F) Fluorescent signals from foreleg neurons across three heat pulse trials, each consisting of three consecutive pulses of heat stimuli. The first (D) and third (F) trials had the Peltier element on, while the second was turned off (E). n = 47 ROI.

**Supplemental Figure S4.**
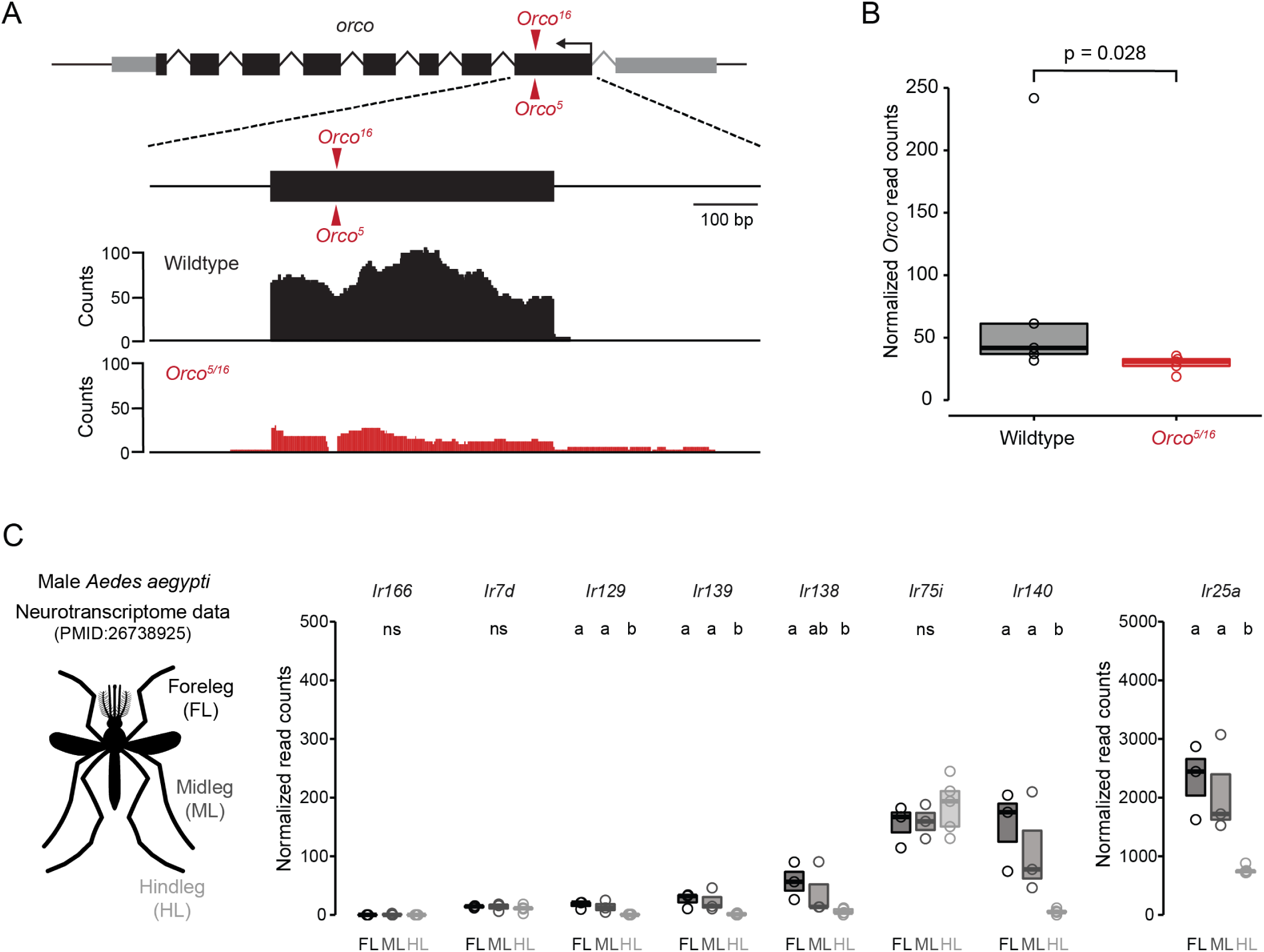
Related to Figure 7 *Aedes aegypti* leg transcriptome analysis. (A) Graphical representation of the number of raw mapped reads at the first exon at the *Orco* locus from wildtype (black) and *Orco^5/16^* mutant leg RNAseq dataset (red). *Orco^5^* and *Orco^16^* loci are indicated with red arrowheads. (B) Normalized *Orco* read counts from wildtype and *Orco^5/16^* legs (DESeq2, FDR < 0.05). (C) Normalized read counts of *Orco^5/16^* leg upregulated IR genes in the male leg neurotranscriptome dataset (*29*). Data labeled with different letters are significantly different (n=3-5 replicates/leg, FDR < 0.01, pairwise DESeq2 comparison). Data are plotted as scatter-box plots (individual data points, median as horizontal line, interquartile range as box) for (B, C).

**Supplemental Figure S5.**
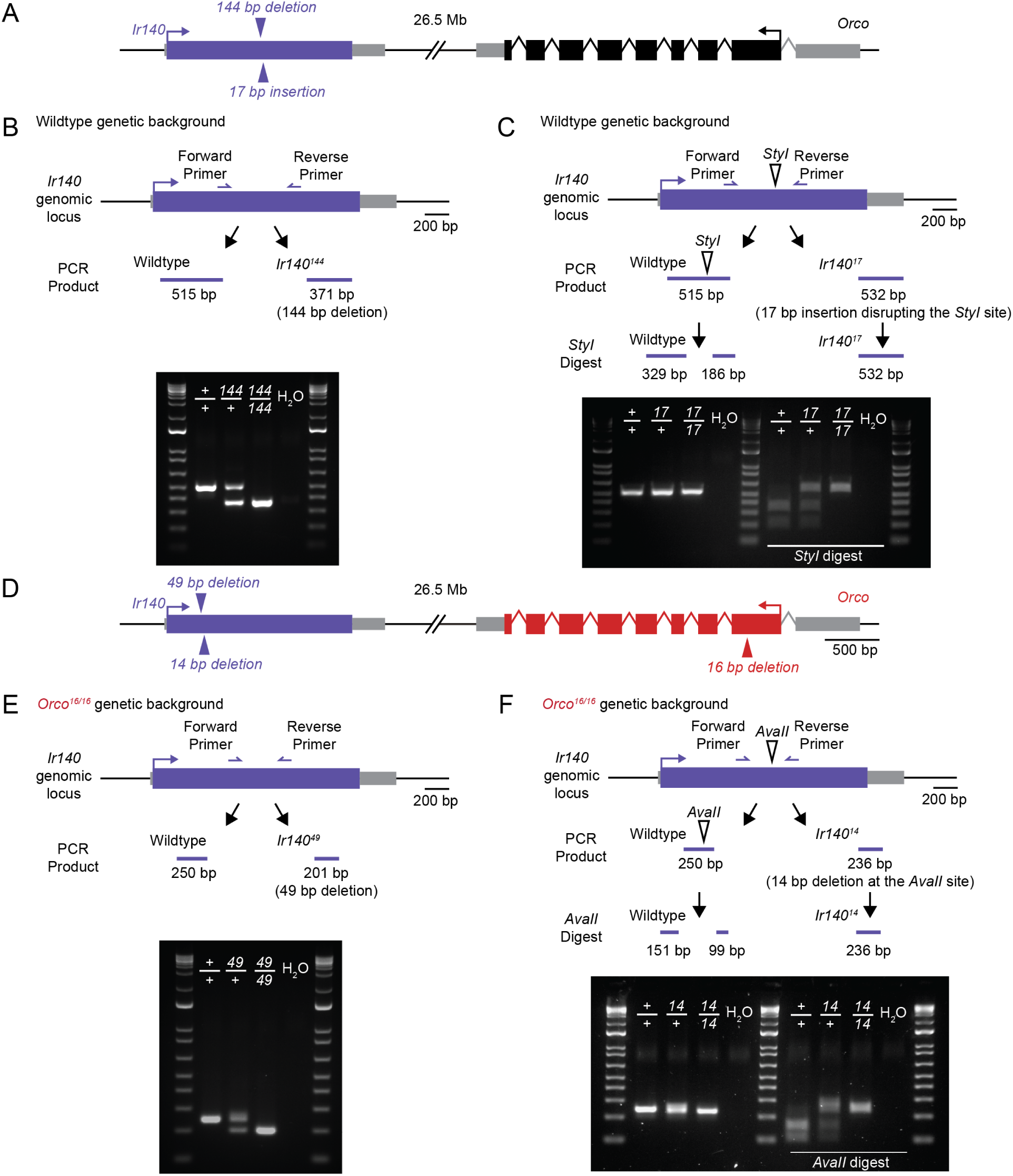
Related to Figure 8 PCR genotyping *Ir140* single and *Orco, Ir140* double mutant strains. (A,D) Schematic of the *Ir140*, *Orco* loci, and the isolated alleles for single (A) and double mutants (D). (B,C,E,F) Genotyping schematics (top) and agarose gel electrophoresis images (bottom) of PCR fragments. PCR fragments before (left) and after (right) restriction digests with the indicated restriction enzymes to genotype the indicated *Ir140* (C) and *Orco, Ir140* double mutants (F).

## SUPPLEMENTAL VIDEOS

**Supplemental Video S1** – Related to Figure 1

**Mosquito heat-seeking behavior**

Percent mosquitoes on Peltier (top left) quantified using images (right) acquired across the 36°C heat stimulus (bottom left). CO_2_ pulse is indicated in green, and the Peltier location is outlined with a dotted line. The images are acquired at 1 Hz, and the movie is sped up 20X.

**Supplemental Video S2** – Related to Figure 2

***Orco^5/16^* heat-seeking behavior**

Heat-seeking behavior comparing wildtype control (middle) and *Orco^5/16^* mutants (right) during the 36°C heat stimulus (left). CO_2_ pulse is indicated in green, and the Peltier location is outlined with a dotted line. The images are acquired at 1 Hz, and the movie is sped up 20X.

**Supplemental Video S3** – Related to Figure 5

**Tarsal neuron distribution along the cranial-caudal axis.**

3D reconstruction of dTomato confocal image of forelegs taken from a *brp-QF2w > QUAS-dTomato-T2A-GCaMP6s* animal.

**Supplemental Video S4** – Related to Figure 6

**Ratiometric calcium imaging of foreleg tarsal neurons.**

An example *ex vivo* foreleg tarsal imaging experiment with three consecutive heat pulses. The right panel is a movie of a ratiometric image of the three most distal tarsal segments. The foreleg is outlined with a white dotted line. The top right panel shows the ratiometric fluorescent signals across the experiment. Each trace indicates ratiometric calcium responses from regions highlighted in the same color as the right panel. The blue trace represents a non-responding cell. The bottom right panel shows the temperature traces. White arrowheads indicate heat pulses.

## REFERENCES

1. N. Lessard, M. Paré, F. Lepore, M. Lassonde, Early-blind human subjects localize sound sources better than sighted subjects. Nature 395, 278–280 (1998).

2. D. Goldreich, I. M. Kanics, Tactile acuity is enhanced in blindness. J Neurosci 23, 3439–3445 (2003).

3. D. Bavelier, H. J. Neville, Cross-modal plasticity: where and how? Nat Rev Neurosci 3, 443–452 (2002).

4. M. Sur, P. E. Garraghty, A. W. Roe, Experimentally induced visual projections into auditory thalamus and cortex. Science 242, 1437–1441 (1988).

5. A. W. Roe, S. L. Pallas, J. O. Hahm, M. Sur, A map of visual space induced in primary auditory cortex. Science 250, 818–820 (1990).

6. A. W. Roe, S. L. Pallas, Y. H. Kwon, M. Sur, Visual projections routed to the auditory pathway in ferrets: receptive fields of visual neurons in primary auditory cortex. J Neurosci 12, 3651–3664 (1992).

7. N. Sadato et al., Activation of the primary visual cortex by Braille reading in blind subjects. Nature 380, 526–528 (1996).

8. R. Weeks et al., A positron emission tomographic study of auditory localization in the congenitally blind. J Neurosci 20, 2664–2672 (2000).

9. I. Rabinowitch et al., Neuropeptide-driven cross-modal plasticity following sensory loss in *Caenorhabditis elegans*. PLoS Biol 14, e1002348 (2016).

10. A. Goel et al., Cross-modal regulation of synaptic AMPA receptors in primary sensory cortices by visual experience. Nat Neurosci 9, 1001–1003 (2006).

11. S. Jitsuki et al., Serotonin mediates cross-modal reorganization of cortical circuits. Neuron 69, 780–792 (2011).

12. P. Junca, M. Stanley, P.-Y. Musso, M. D. Gordon, Modulation of taste sensitivity by the olfactory system in *Drosophila*. bioRxiv, 10.1101/2021.1103.1130.437740 (2021).

13. C. J. McMeniman, R. A. Corfas, B. J. Matthews, S. A. Ritchie, L. B. Vosshall, Multimodal integration of carbon dioxide and other sensory cues drives mosquito attraction to humans. Cell 156, 1060–1071 (2014).

14. W. D. Jones, P. Cayirlioglu, I. G. Kadow, L. B. Vosshall, Two chemosensory receptors together mediate carbon dioxide detection in *Drosophila*. Nature 445, 86–90 (2007).

15. J. Y. Kwon, A. Dahanukar, L. A. Weiss, J. R. Carlson, The molecular basis of CO_2_ reception in *Drosophila*. Proc Natl Acad Sci U S A 104, 3574–3578 (2007).

16. L. Abuin et al., Functional architecture of olfactory ionotropic glutamate receptors. Neuron 69, 44–60 (2011).

17. R. Benton, S. Sachse, S. W. Michnick, L. B. Vosshall, Atypical membrane topology and heteromeric function of *Drosophila* odorant receptors in vivo. PLoS Biol 4, e20 (2006).

18. B. J. Matthews et al., Improved reference genome of *Aedes aegypti* informs arbovirus vector control. Nature 563, 501–507 (2018).

19. L. Ni, The structure and function of Ionotropic Receptors in *Drosophila*. Front Mol Neurosci 13, 638839 (2020).

20. M. DeGennaro et al., *orco* mutant mosquitoes lose strong preference for humans and are not repelled by volatile DEET. Nature 498, 487–491 (2013).

21. K. E. Kistler, L. B. Vosshall, B. J. Matthews, Genome engineering with CRISPR-Cas9 in the mosquito *Aedes aegypti*. Cell Rep 11, 51–60 (2015).

22. J. I. Raji et al., *Aedes aegypti* mosquitoes detect acidic volatiles found in human odor using the *IR8a* pathway. Curr Biol 29, 1253–1262 e1257 (2019).

23. Z. Ye et al., Discrete roles of Ir76b ionotropic coreceptor impact olfaction, blood feeding, and mating in the malaria vector mosquito *Anopheles coluzzii*. Proc Natl Acad Sci U S A 119, e2112385119 (2022).

24. C. Greppi et al., Mosquito heat seeking is driven by an ancestral cooling receptor. Science 367, 681–684 (2020).

25. W. J. Laursen et al., Humidity sensors that alert mosquitoes to nearby hosts and egg-laying sites. Neuron 111, 874–887 e878 (2023).

26. M. Herre et al., Non-canonical odor coding in the mosquito. Cell 185, 3104–3123.e3128 (2022).

27. D. Task et al., Chemoreceptor co-expression in *Drosophila melanogaster* olfactory neurons. Elife 11, (2022).

28. R. A. Corfas, L. B. Vosshall, The cation channel TRPA1 tunes mosquito thermotaxis to host temperatures. Elife 4, (2015).

29. B. J. Matthews, C. S. McBride, M. DeGennaro, O. Despo, L. B. Vosshall, The neurotranscriptome of the *Aedes aegypti* mosquito. BMC Genomics 17, 32 (2016).

30. R. Benton, K. S. Vannice, C. Gomez-Diaz, L. B. Vosshall, Variant ionotropic glutamate receptors as chemosensory receptors in *Drosophila*. Cell 136, 149–162 (2009).

31. C. Chen et al., *Drosophila* Ionotropic Receptor 25a mediates circadian clock resetting by temperature. Nature 527, 516–520 (2015).

32. A. Enjin et al., Humidity sensing in *Drosophila*. Curr Biol 26, 1352–1358 (2016).

33. Z. A. Knecht et al., Ionotropic Receptor-dependent moist and dry cells control hygrosensation in *Drosophila*. Elife 6, (2017).

34. G. Budelli et al., Ionotropic receptors specify the morphogenesis of phasic sensors controlling rapid thermal preference in *Drosophila*. Neuron 101, 738–747 e733 (2019).

35. M. E. De Obaldia et al., Differential mosquito attraction to humans is associated with skin-derived carboxylic acid levels. Cell 185, 4099–4116 e4013 (2022).

36. Y. V. Zhang, J. Ni, C. Montell, The molecular basis for attractive salt-taste coding in *Drosophila*. Science 340, 1334–1338 (2013).

37. A. Ganguly et al., A molecular and cellular context-dependent role for Ir76b in detection of amino acid taste. Cell Rep 18, 737–750 (2017).

38. Y. Chen, H. Amrein, Ionotropic receptors mediate *Drosophila* oviposition preference through sour gustatory receptor neurons. Curr Biol 27, 2741–2750 e2744 (2017).

39. J. E. Ahn, Y. Chen, H. Amrein, Molecular basis of fatty acid taste in *Drosophila*. Elife 6, (2017).

40. M. Ai et al., Ionotropic glutamate receptors IR64a and IR8a form a functional odorant receptor complex in vivo in *Drosophila*. J Neurosci 33, 10741–10749 (2013).

41. E. J. Dennis, O. V. Goldman, L. B. Vosshall, *Aedes aegypti* mosquitoes use their legs to sense DEET on contact. Curr Biol 29, 1551–1556 (2019).

42. L. Ni et al., A gustatory receptor paralogue controls rapid warmth avoidance in *Drosophila*. Nature 500, 580–584 (2013).

43. J. M. Simoes et al., Robustness and plasticity in *Drosophila* heat avoidance. Nat Commun 12, 2044 (2021).

44. M. Gallio, T. A. Ofstad, L. J. Macpherson, J. W. Wang, C. S. Zuker, The coding of temperature in the *Drosophila* brain. Cell 144, 614–624 (2011).

45. F. N. Hamada et al., An internal thermal sensor controlling temperature preference in *Drosophila*. Nature 454, 217–220 (2008).

46. J. Liu et al., Alleviation of thermal nociception depends on heat-sensitive neurons and a TRP channel in the brain. Curr Biol 33, 2397–2406 e2396 (2023).

47. E. E. Davis, P. G. Sokolove, Temperature responses of antennal receptors of the mosquito, *Aedes aegypti*. J. Comp. Physiol. A 96, 223–236 (1975).

48. E. Gingl, A. Hinterwirth, H. Tichy, Sensory representation of temperature in mosquito warm and cold cells. J Neurophysiol 94, 176–185 (2005).

49. G. Wang et al., *Anopheles gambiae* TRPA1 is a heat-activated channel expressed in thermosensitive sensilla of female antennae. Eur J Neurosci 30, 967–974 (2009).

50. L. M. Roth, Loci of sensory end-organs used by mosquitoes (*Aedes aegypti* (L.) and Anopheles quadrimaculatus say) in receiving host stimuli. Ann Entomol Soc Am 44, 59–74 (1951).

51. Y. Zhou, D. Deng, R. Chen, C. Lai, Q. Chen, Effects of antennal segments defects on blood-sucking behavior in *Aedes albopictus*. PLoS One 18, e0276036 (2023).

52. A. L. Costa-da-Silva et al., Glytube: a conical tube and parafilm M-based method as a simplified device to artificially blood-feed the dengue vector mosquito, *Aedes aegypti*. PLoS One 8, e53816 (2013).

53. T. R. Sorrells, A. Pandey, A. Rosas-Villegas, L. B. Vosshall, A persistent behavioral state enables sustained predation of humans by mosquitoes. Elife 11, (2022).

54. A. L. Carr, V. L. Salgado, Ticks home in on body heat: A new understanding of Haller’s organ and repellent action. PLoS One 14, e0221659 (2019).

55. F. Ling, A. Dahanukar, L. A. Weiss, J. Y. Kwon, J. R. Carlson, The molecular and cellular basis of taste coding in the legs of *Drosophila*. J Neurosci 34, 7148–7164 (2014).

56. E. G. Freeman, A. Dahanukar, Molecular neurobiology of *Drosophila* taste. Curr Opin Neurobiol 34, 140–148 (2015).

57. Z. Zhao, D. Tian, C. S. McBride, Development of a pan-neuronal genetic driver in *Aedes aegypti* mosquitoes. Cell Rep Methods 1, (2021).

58. V. Jové et al., Sensory discrimination of blood and floral nectar by *Aedes aegypti* mosquitoes. Neuron 108, 1163–1180 (2020).

59. R. Xiao et al., A genetic program promotes *C. elegans* longevity at cold temperatures via a thermosensitive TRP channel. Cell 152, 806–817 (2013).

60. J. Gong et al., A cold-sensing receptor encoded by a glutamate receptor gene. Cell 178, 1375–1386 e1311 (2019).

61. W. D. Tracey, R. I. Wilson, G. Laurent, S. Benzer, *painless*, a *Drosophila* gene essential for nociception. Cell 113, 261–273 (2003).

62. S. Y. Xu et al., Thermal nociception in adult *Drosophila*: behavioral characterization and the role of the painless gene. Genes Brain Behav 5, 602–613 (2006).

63. Y. Lee et al., Pyrexia is a new thermal transient receptor potential channel endowing tolerance to high temperatures in *Drosophila melanogaster*. Nat Genet 37, 305–310 (2005).

64. M. Rosenzweig et al., The *Drosophila* ortholog of vertebrate *TRPA1* regulates thermotaxis. Genes Dev 19, 419–424 (2005).

65. L. Ni et al., The Ionotropic Receptors IR21a and IR25a mediate cool sensing in *Drosophila*. Elife 5, (2016).

66. Y. Kwon, W. L. Shen, H. S. Shim, C. Montell, Fine thermotactic discrimination between the optimal and slightly cooler temperatures via a TRPV channel in chordotonal neurons. J Neurosci 30, 10465–10471 (2010).

67. W. L. Shen et al., Function of rhodopsin in temperature discrimination in *Drosophila*. Science 331, 1333–1336 (2011).

68. T. Sokabe, H. C. Chen, J. Luo, C. Montell, A switch in thermal preference in *Drosophila* larvae depends on multiple rhodopsins. Cell Rep 17, 336–344 (2016).

69. R. Thistle, P. Cameron, A. Ghorayshi, L. Dennison, K. Scott, Contact chemoreceptors mediate male-male repulsion and male-female attraction during *Drosophila* courtship. Cell 149, 1140–1151 (2012).

70. B. N. A. Hudson, The behaviour of the female mosquito in selecting water for oviposition. J Exp Biol 33, 478–492 (1956).

71. B. J. Matthews, M. A. Younger, L. B. Vosshall, The ion channel *ppk301* controls freshwater egg-laying in the mosquito *Aedes aegypti*. Elife 8, e43963 (2019).

72. M. Z. Liu, L. B. Vosshall, General visual and contingent thermal cues interact to elicit attraction in female *Aedes aegypti* mosquitoes. Curr Biol 29, 2250–2257 e2254 (2019).

73. O. S. Akbari et al., The developmental transcriptome of the mosquito *Aedes aegypti*, an invasive species and major arbovirus vector. G3 (Bethesda) 3, 1493-1509 (2013).

74. O. G. David et al., Fertility decline in female mosquitoes is regulated by the orco olfactory co-receptor. *bioRxiv*, 10.1101/2023.1101.1107.523099 (2023).

75. N. S. Basrur et al., *Fruitless* mutant male mosquitoes gain attraction to human odor. Elife 9, e63982 (2020).

76. S. N. Fry, M. Bichsel, P. Müller, D. Robert, Tracking of flying insects using pan-tilt cameras. J Neurosci Methods 101, 59–67 (2000).

77. R. Patro, G. Duggal, M. I. Love, R. A. Irizarry, C. Kingsford, Salmon provides fast and bias-aware quantification of transcript expression. Nat Methods 14, 417–419 (2017).

78. C. Soneson, M. I. Love, M. D. Robinson, Differential analyses for RNA-seq: transcript-level estimates improve gene-level inferences. F1000R es 4, 1521 (2015).

79. M. I. Love, W. Huber, S. Anders, Moderated estimation of fold change and dispersion for RNA-seq data with DESeq2. Genome Biol 15, 550 (2014).

80. A. Dobin et al., STAR: ultrafast universal RNA-seq aligner. Bioinformatics 29, 15–21 (2013).

81. H. Thorvaldsdottir, J. T. Robinson, J. P. Mesirov, Integrative Genomics Viewer (IGV): high-performance genomics data visualization and exploration. Brief Bioinform 14, 178–192 (2013).

82. M. Ohkura et al., Genetically encoded green fluorescent Ca2+ indicators with improved detectability for neuronal Ca2+ signals. PLoS One 7, e51286 (2012).

83. N. Helassa, B. Podor, A. Fine, K. Török, Design and mechanistic insight into ultrafast calcium indicators for monitoring intracellular calcium dynamics. Sci Rep 6, 38276 (2016).

84. K. Labun et al., CHOPCHOP v3: expanding the CRISPR web toolbox beyond genome editing. Nucleic Acids Res 47, W171–W174 (2019).

85. N. Jasinskiene, J. Juhn, A. A. James, Microinjection of *A. aegypti* embryos to obtain transgenic mosquitoes. J Vis Exp, 219 (2007).

86. RCoreTeam, R: A language and environment for statistical computing. *R Foundation for Statistical Computing, Vienna, Austria.* https://www.R-project.org/. (2021).

